# Extensive introgression among North American wild grapes (*Vitis*) fuels biotic and abiotic adaptation

**DOI:** 10.1101/2021.02.11.430822

**Authors:** Abraham Morales-Cruz, Jonas Aguirre-Liguori, Yongfeng Zhou, Andrea Minio, Summaira Riaz, Andrew M. Walker, Dario Cantu, Brandon S. Gaut

## Abstract

Introgressive hybridization can introduce adaptive genetic variation into a species or population. To evaluate the evolutionary forces that contribute to introgression, we studied six *Vitis* species that are native to the Southwestern United States and potentially useful for breeding grapevine (*V. vinifera*) rootstocks. By creating a reference genome from one wild species, *V. arizonica*, and by resequencing 130 accessions, we focused on identifying putatively introgressed regions (pIRs) between species. We found that up to ~8% of extant genome is attributable to introgression between species. The pIRs tended to be gene poor, located in regions of high recombination and enriched for genes implicated in disease resistance functions. To assess potential pIR function, we explored SNP associations to bioclimatic variables and to bacterial levels after infection with the causative agent of Pierce’s Disease. pIRs were enriched for SNPs associated with both climate and bacterial levels, suggesting potential drivers of adaptive events. Altogether, this study yields insights into the genomic extent of introgression, potential pressures that shape adaptive introgression, and the history of economically important wild relatives of a critical crop.

## INTRODUCTION

Species emerge from complex interactions among evolutionary processes. For example, genetic drift and local adaptation drive divergence between populations, which ultimately leads to genetic isolation and eventual speciation (Rundle & Nosil, 2005; Wu, 2001). The process of divergence can be slowed, in turn, by gene flow between populations, which maintains genetic similarity. There is growing evidence, however, that introgressive hybridization between populations and species does more than homogenize gene pools, because it may also be a source of novelty that reassorts genetic variants into beneficial combinations, permitting adaptation to new ecological niches (Marques et al., 2019). This ability to reassort genetic variation may partially explain the highly reticulated evolutionary history of adaptive radiations like *Heliconius* butterflies (Edelman et al., 2019), tomatoes (Pease et al., 2016), Darwin’s finches (Lamichhaney et al., 2015) and African cichlids (Meier et al., 2017). Introgression has also played a major role in the diversification and speciation of angiosperms (Anderson & Stebbins, 1954), affecting an estimated ~25% of flowering plant species (Mallet, 2005).

It is generally not known how frequently introgression occurs between species, whether introgression events are adaptive, and, if so, the traits that have been affected. Fortunately, genomic approaches have begun to provide some insights into these central questions. For example, the analysis of *Heliconius* genomes suggests that a large inversion was transferred between species and that this event was adaptive because the inversion contains a color pattern locus that controls mimicry and crypsis (Edelman et al., 2019). Similarly, a large chromosomal region was exchanged between distinct sunflower subspecies, likely facilitating genetic adaptation to xeric environments (Todesco et al., 2020). Recent work in cypress (Y. Ma et al., 2019), oaks (Leroy et al., 2020; Nagamitsu et al., 2020), maize (Hufford et al., 2013) and cultivated date palms (Flowers et al., 2019) also suggest that introgression between plant species facilitates adaptation to local environments, although in most cases the phenotypic basis for adaptation is not clear. The work in maize and date palms further highlights the importance of studying the wild relatives of crop species, because they are potentially important sources of adaptive traits for agronomic improvement (Burgarella et al., 2019).

Here we study introgression among members of the genus *Vitis*, a genus of ~70 species that have radiated to varied environments. The genus likely originated in North America ~ 45my (Z.-Y. Ma, Wen, Ickert-Bond, et al., 2018) and contains two subgenera: *Muscadinia* (2*n*=40) and *Vitis* (2*n*=38). The subgenus *Vitis* has a disjunct distribution across North America and Eurasia (Z.-Y. Ma, Wen, Ickert-Bond, et al., 2018), with the ~25 North American species (Wan et al., 2013) distributed broadly across the continent, including the American Southwest, where extreme temperature changes and drought are pervasive abiotic stressors. All species within the subgenus are dioecious, interfertile, and often sympatric (Heinitz et al., 2019), suggesting the possibility of an extensive history of introgression among species (Aradhya et al., 2013; Z.-Y. Ma, Wen, Tian, et al., 2018). However, the extent and genomic location of introgression regions remains unexplored, as do the potential functions and evolutionary forces that led to the retention of those regions.

*Vitis* is also an important study system because of its economic value. The domesticated grapevine (*Vitis vinifera* spp. *vinifera;* hereafter *vinifera*) is arguably the most valuable horticultural crop in the world (Alston & Sambucci, 2019) and an excellent model for the study of perennial fruit crops (Minio et al., 2017). It is not always appreciated, however, that the cultivation, sustainability, and security of grapevine cultivation relies on North American (NA) *Vitis* species as sources of resistance to abiotic and biotic stresses (Heinitz et al., 2019; Walker et al., 2014; Warschefsky et al., 2016). The susceptibility of cultivated *vinifera* was apparent after its introduction to North America in the late 1600s, because grapevines quickly succumbed to diseases like powdery mildew (*Erysiphe necator* Schwein.), downy mildew (*Plasmopara viticola* (Berk & Curt.) Berl. & de Toni), phylloxera (*Daktulosphaira vitifoliae* Fitch) and abiotic stresses such as cold (Heinitz et al., 2019). The recognition of *vinifera* susceptibility, especially to phylloxera, and the later unintended dissemination of these pests and diseases into Europe, launched breeding efforts to use NA *Vitis* as rootstocks (Heinitz et al., 2019; Summaira Riaz et al., 2019). The use of NA *Vitis* as rootstocks has been so successful that an estimated 80% of viticulture across the globe currently relies on grafting (Ollat et al., 2016). Currently, about a dozen NA *Vitis* species are actively utilized for breeding (This et al., 2006), but the few major rootstock cultivars currently in use represent a narrow genetic background (Summaira Riaz et al., 2019).

Despite the successful integration of NA *Vitis* into viticulture, there is ongoing pressure to identify additional sources of resistance to biotic and abiotic stress. As an example of the former, Pierce’s Disease (PD) has been a problematic disease of grapevines, and it is currently considered a global threat to the sustainability of wine production (Purcell & Saunders, 1999). PD is caused by a bacterium (*Xyllela fastidiosa*) that spreads from plant to plant by xylem-feeding insect vectors. Increased disease pressure from PD is caused in part by shifts in the range of these insect vectors due to climate change. Interestingly, several southeastern *Vitis* species and muscadine grapes (*M. rotundofolia*) show resistance to PD, but the best-studied PD resistant locus (*PdR1*) was characterized from *V. arizonica* (Krivanek et al., 2006), a native to the Southwestern United States. A subsequent study has shown that *PdR1* is also present in other wild species (S. Riaz et al., 2018) and that it is not the only PD resistance locus (S. Riaz et al., 2018). These observations open interesting questions about the evolutionary history of PD resistance, and more generally about the potential introgression of pathogen resistance loci among wild grape species.

In this study, we focus on the extent and pattern of introgression among *Vitis* species. To do so, we have generated whole-genome resequencing data from 130 accessions representing six *Vitis* species from portions of their native ranges in eight Southwestern U.S. states (**Figure 1, Figure S1**). Some of these species have largely overlapping distributions (e.g., *V candicans* and *V. berlandieri* in Texas), others have disjunct distributions (e.g., *V. arizonica*) and still another (*V. riparia*) has populations in the Southwest at the edge of a broader continental distribution. We have chosen to focus on these *Vitis* from the Southwest because: *i*) they are potentially useful as breeding material for resistance to abiotic stresses (Heinitz et al., 2019), *ii*) several have been utilized in rootstock breeding already (Doucleff et al., 2004; This et al., 2006), and *iii*) they are polymorphic for tolerance and susceptibility to PD (Krivanek et al., 2006; S. Riaz et al., 2008; Summaira Riaz et al., 2020). To complement genetic data, we have also assessed PD resistance for each accession and gathered bioclimatic data from their location of origin. Given this multifaceted dataset, we address three sets of questions. First, given that species are interfertile and can overlap substantively in geographic range, are they genetically distinct? Second, if they are distinct, is there nonetheless genetic evidence for introgression? Third, if there is evidence for introgression, what are the genomic characteristics of introgressed regions, in terms of locations, size and gene content? Finally, is there evidence that introgression events have played an adaptive role, as evidenced by genetic associations with either disease resistance or bioclimatic variables?

**Figure 1.**
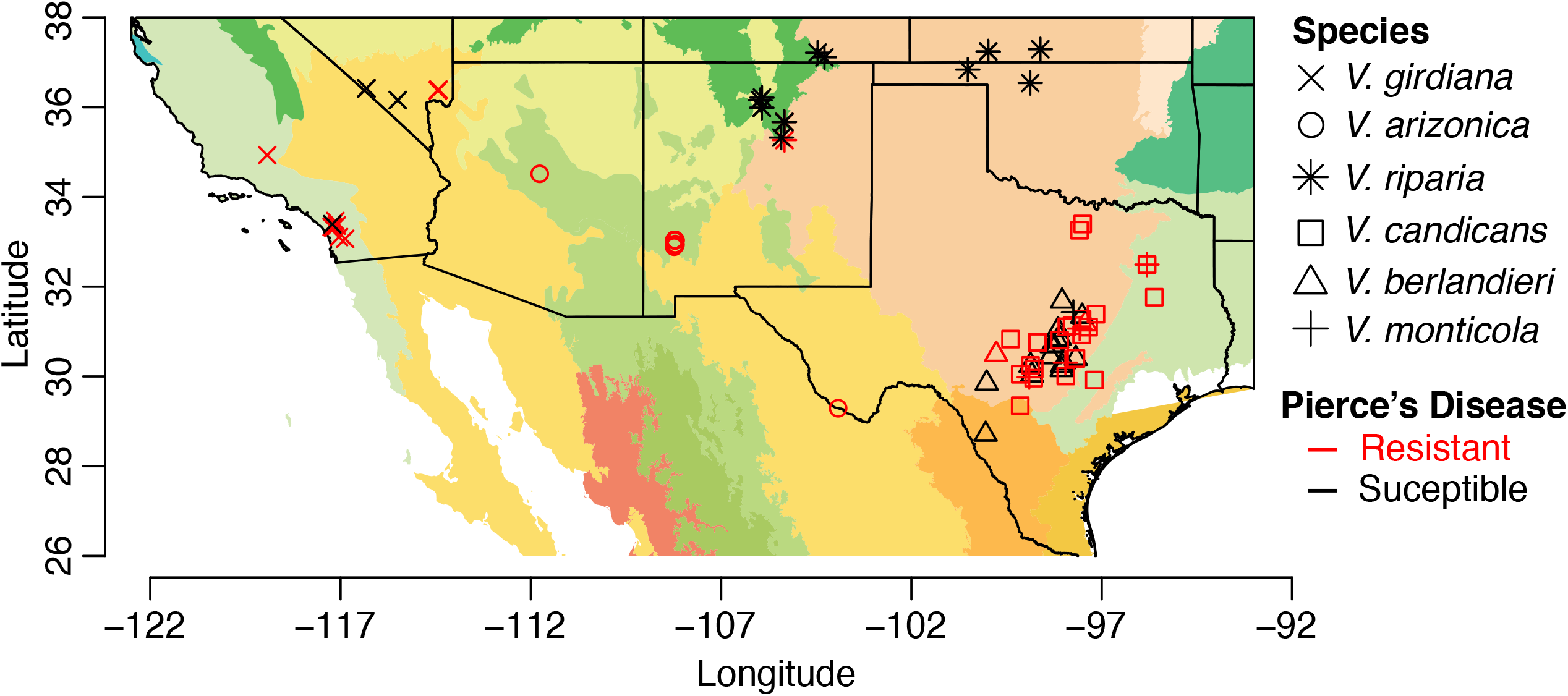
Geographic distribution and ecological diversity of sampled populations of wild grapes. Shapes correspond to different species, and samples colored red or black were classified as resistant or susceptible to Pierce’s Disease, respectively. The colors of the regions correspond to the level II ecological region as defined by the United States Environmental Protection Agency (EPA). See Figure S1 for additional details.

## RESULTS

### Population structure and phylogeny

We resequenced the whole genome of 130 *Vitis* samples from throughout the native range of six species (*V. arizonica, V. berlandieri, V. candicans, V. girdiana, V. monticola*, and *V. riparia*) (**Figure 1, Table S1**). We mapped resequencing data to a reference genome of *V. arizonica* produced from long-read sequences (Massonnet et al., 2020). After assembling sequencing reads with an optical map, the reference assembly contained 19 anchored pseudomolecules, an N50 of 25.9 Mb, a size of 503 Mb and a BUSCO score of 96.4% (**Table S2 & S3**). We used long-read RNAseq data and *de novo* analyses to annotate genes with the genome, ultimately predicting 28,259 gene models.

After mapping resequencing data to the reference, we identified ~ 20 million SNPs among all samples (**Table S4**) and used them to assess genetic structure using NGSadmix (Skotte et al., 2013) with *K=1* to 10 clusters. The highest support was for *K=7* clusters, corresponding to one per species, except for *V. girdiana*, which had two disjunct groups from different geographical locations (**Figure 1 & 2A, Figure S2**). Based on the results, we found and removed 19 hybrid individuals that had <80% of the admixture proportion assigned to a single cluster (Castillo et al., 2010; Vigouroux et al., 2008), leaving a final dataset of 111 accessions. The remaining 111 samples included accessions from *V. arizonica* (n= 22), *V. candicans* (n= 24), *V. berlandieri* (n= 22), *V. girdiana* (n= 18), *V. riparia* (n= 19) and *V. monticola* (*n*= 6) (**Figure 2A**). We calculated genome-wideπfor each species; it averaged 0.00284 across species and ranged from 0.00211 in *V. berlandieri* to 0.00353 in *V. monticola* (**Table S5**).

**Figure 2.**
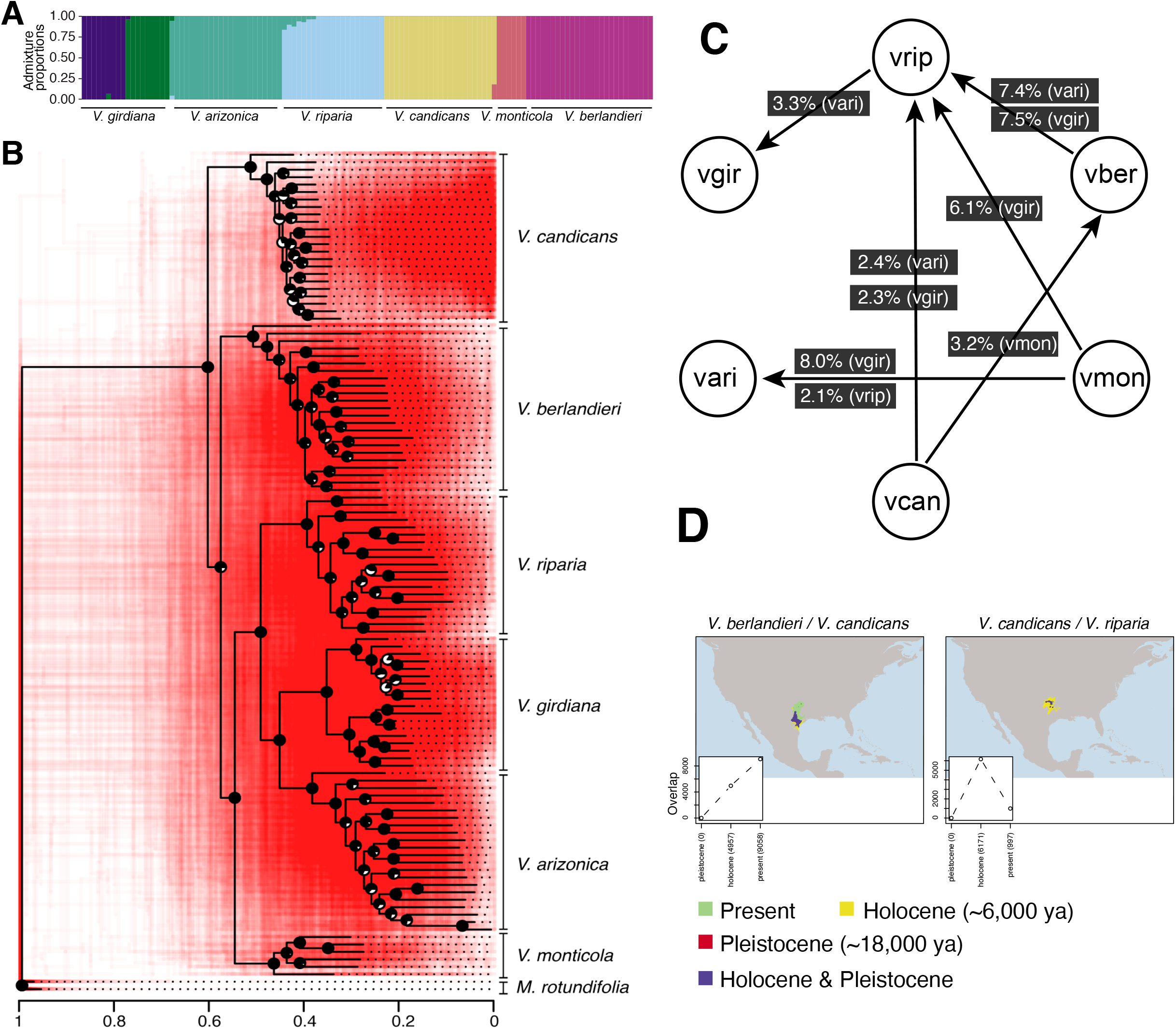
Genetic history of the wild grapes sampled. (A) Genetic structure of samples with colors representing each cluster detected by the structure analysis (K=7). Hybrid samples are not included but see Figure S2. (B) The phylogenetic tree in black corresponds to the consensus tree. Each node has a pie chart with the black portion indicating the proportion of supporting bootstrap replicates. The red phylogenies in the background correspond to 500 highly supported consensus trees (median bootstrap support > 70%) based on separate 10 Kb windows throughout the genome. (C) Diagram summarizing significant signals of introgression. Arrows show the direction of the introgression, from donor to the recipient species. The numbers next to the arrows are the overall f4-ratio value per trio, and the abbreviation in the black box is the “control” population used in the test. In this diagram, the species are abbreviated as mrot: M. rotundifolia, vari: *V. arizonica, vcan: V. candicans, vmon: V. monticola, vber: V. berlandieri, vrip: V. riparia and vgir: V. girdiana.* (D) Examples of SDMs overlaps from pairs of species with evidence of introgression projected in at least one of three periods: the present, the Holocene and the Pleistocene. The inset in the bottom left corner shows the amount of overlap per period. See Figures S4, S5 and S6 for additional SDMs featuring pairs of species.

We created a consensus phylogenetic tree based on a reduced number of SNPs to limit the effects of linkage disequilibrium (see Methods). The phylogeny had median bootstrap support of 88.5% for all nodes and strong support (> 76%) for nodes that separated species (**Figure 2B, Figure S3**). Each species was monophyletic, showing that each species is genetically identifiable and justifying treating each named species as a separate group. In contrast, phylogenies based on individual 10Kb genomic regions from throughout the genome were highly discordant, representing ‘clouds’ around the grouping of species (**Figure 2B**). These clouds suggest the possibility of incomplete lineage sorting or substantive histories of introgression among species.

### Tests for introgression and geographic overlap among species

To formally test for introgression, we calculated the *D* statistic (Durand et al., 2011; Patterson et al., 2012). *D* is a genome-wide statistic that measures the excess of shared ancestral alleles in one of two sister species (a receptor and a control) from a third species (the potential donor); it is expected to be zero under the null hypothesis of lineage sorting, but deviates from zero when there is introgression (Durand et al., 2011; Patterson et al., 2012). We calculated *D* for all combinations of three species (hereafter called ‘trios’) that had an appropriate topology for the test according to the consensus tree (**Figure 2B, Table S6**). We used *Muscadinia rotundifolia* as the outgroup for all trios and estimated significance using a block jackknife approach (Patterson et al., 2012).

Of 11 trios tested, nine had a significant *D* value at *p* < 0.0031 (**Table S6**), representing a total of six donor-receptor pairs (**Figure 2C**). The *D* statistic is useful for detecting introgression but a poor estimator of the introgressed proportion of genome (Milan Malinsky et al., 2020; Martin et al., 2015), so we applied the *f*4-ratio (Patterson et al., 2012) to estimate the proportion of the receptor genome that came from a donor. The values ranged from ~2.3% of the genome derived in *V. riparia* from *V. candicans* to ~8.0% in *V. arizonica* from *V. monticola* (**Figure 2C & Table 1**). For some comparisons, we were able to test the same donor-receptor pairs with different control taxa. In two cases, the same donor-receptor pairs yielded similar estimates - e.g., *V. berlandieri* contributed ~7.5% to *V. riparia* andr *V. candicans* donated ~2.4% to *V. riparia* (**Figure 2C**). In contrast, the *f*4 estimate varied widely, from 2.1% to 8.0%, for *V. monticola* donating to *V. arizonica* depending on the control species; notably, however, *D* was significant with either control. Taken together, these results show that: *i*) *D* and *f*4 support historical introgression among species, *ii*) the history of introgression is complex, with at least two species acting as both receptor and donor and iii) our sample of *V. riparia* was most commonly implicated in introgression events (**Figure 3B**).

**Figure 3.**
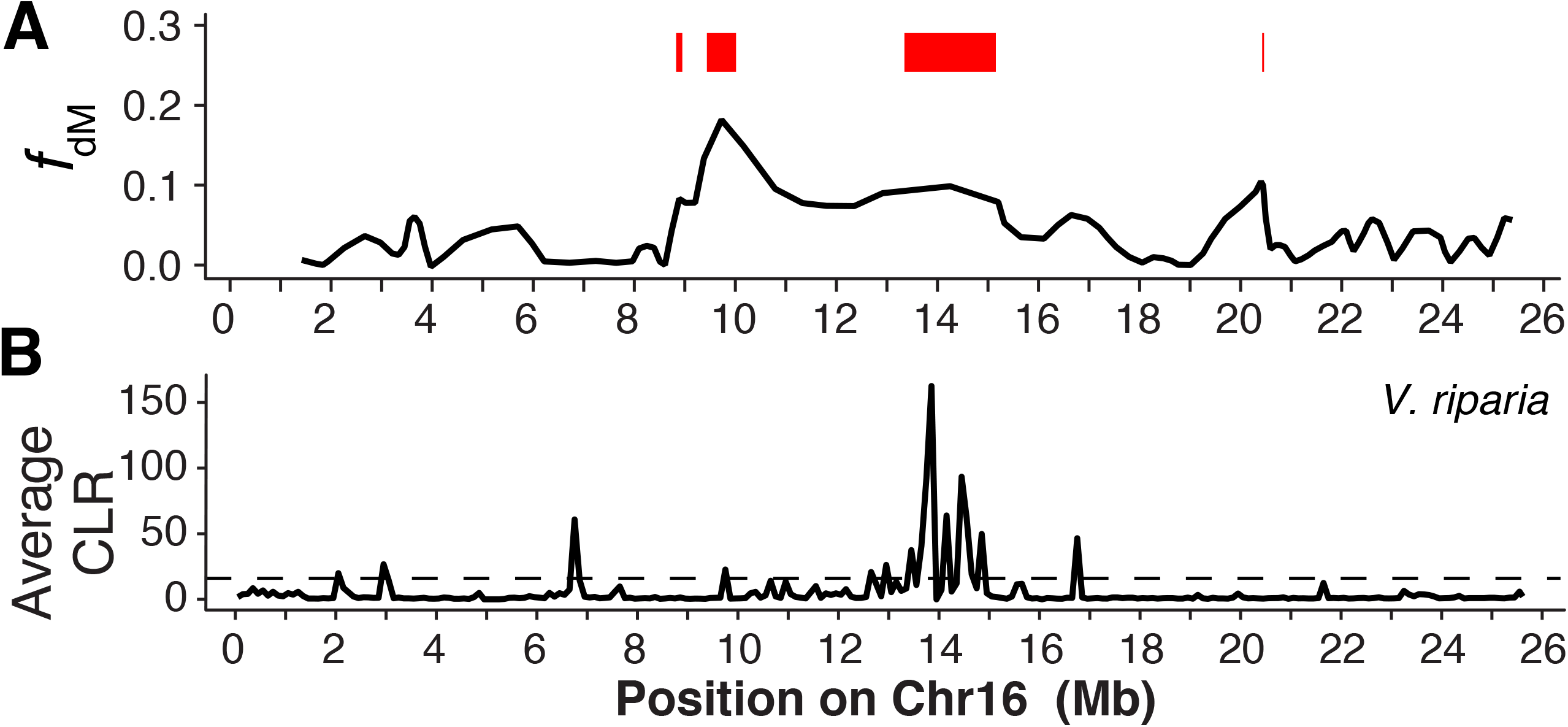
Introgression statistics along chromosome 16 in the ARC trio, which consists of *V. arizonica* as the control, V. riparia as the receptor and *V. candicans* as the donor. (A) Introgression signal measured as fdM in windows across the chromosome. The red regions on top show the pIRs defined by highest x% of fdM values, where x was determined for each receptor species in the trio by the f4 estimate. (B) The CLR metric showing potential locations of selective sweeps in *V. .riparia.*

**Table 1:** Sets of taxa with significant D statistics and properties of pIRs inferred within the receptor species.

| Trio<sup>1</sup> | Receptor Sp. | Donor Sp. | *f*<sup>4</sup> | Mean pIRs (Kb)<sup>2</sup> | No. genes | Gene density<sup>3</sup> | Sweeps<sup>4</sup> | cM/Kb<sup>5</sup> | Resistance genes<sup>6</sup> | PD<sup>7</sup> | Climate<sup>8</sup> |
| --- | --- | --- | --- | --- | --- | --- | --- | --- | --- | --- | --- |
| GAM | *arizonica* | *monticola* | 8.03% | 269.36 | 2282 | 0.96 | 0.27 | 2.27 | 1.1 | 1.25* | 0.32 |
| RAM | *arizonica* | *monticola* | 2.07% | 210.44 | 597 | 0.92 | 0 | 1.1 | 0.56 | 1.57* | 1.25* |
| MBC | *berlandieri* | *candicans* | 3.24% | 163.23 | 746 | 0.82* | 0.39 | 1.52 | 1.12 | 1.97* | 1.99* |
| AGR | *girdiana* | *riparia* | 3.32% | 154.49 | 955 | 0.99 | 0.32 | 1.61 | 0.52 | 0.00 | 1.49* |
| GRM | *riparia* | *monticola* | 6.08% | 270.68 | 1325 | 0.79* | 0.57 | 1.21 | 1.56* | 0.99 | 0.42 |
| GRB | *riparia* | *berlandieri* | 7.47% | 233.29 | 1538 | 0.75* | 0.93 | 1.43 | 2.00* | 0.94 | 1.12* |
| ARB | *riparia* | *berlandieri* | 7.40% | 200.07 | 1607 | 0.75* | 0.77 | 1.8 | 1.73* | 1.55* | 1.27* |
| ARC | *riparia* | *candicans* | 2.43% | 226.25 | 480 | 0.77* | 1.41* | 1.23 | 3.55* | 0.46 | 1.12* |
| GRC | *riparia* | *candicans* | 2.30% | 217.32 | 455 | 0.76* | 0.43 | 1.45 | 3.01* | 0.49 | 1.36* |

<sup>1</sup>The three letters in a trio represent three species that act as the control, receptor and donor, respectively, in introgression tests. The species are A: *V. arizonica*, B: *V. berlandieri*, C: *V. candicans*, G: *V. girdiana*, M: *V. monticola* and R: *V. riparia*. For each trio, *M. rotundifolia* was used as the outgroup for introgression tests.
<sup>2</sup>After merging windows separated by < 1kb in distance
<sup>3</sup>Gene density inside pIRs relative to the average gene density of the whole genome. Asterisks denote trios with significantly lower values of gene density than the whole genome.
<sup>4</sup>Sweeps refers to the relative enrichment of SNPs in pIR regions of the receptor species. Significant enrichment is denoted by an asterisk, with significance assessed by permutation.
<sup>5</sup>cM/kb refers to the ratio of the average recombination rate within pIR regions of the receptor species, relative to the genomic background.
<sup>6</sup>Resistance genes refers to the enrichment of annotated disease-resistance genes in the receptor.
<sup>7</sup>PD refers to the enrichment of Pierce's Disease associated SNPs within pIRs. Significant enrichment is denoted by an asterisk, with significance assessed by permutation.
<sup>8</sup>Climate refers to the enrichment of Climate associated SNPs within pIRs. Significant enrichment is denoted by an asterisk, with significance assessed by permutation.

A puzzling feature of these results is that some receptor-donor pairs have few or no regions of geographic overlap in the present (**Figure 2D; Figure S4**), suggesting limited opportunities for hybridization. To better explore the possibility of sympatry between species, we performed species distribution modeling (SDM). SDMs identify climatic factors that define the geographic distribution of a species, and they then predict the change in species’ distribution over time, given climate prediction models (Fourcade et al., 2017; Peterson et al., 2011; Thullier et al., 2019). We constructed SDMs based on bioclimatic data from the present, the Holocene (~6,000 ya), and the Pleistocene (~18,000 ya) (see Methods). Some of the species pairs had no predicted geographic overlap in any of the three periods (**Figure S5**), and none of these yielded genetic evidence for introgression. Others had overlapping distributions but without detected introgression events (**Figure S6**). Finally, all species-pairs with genomic evidence for introgression had some geographic overlap (**Figure S4**). In some cases, the predicted geographic overlap between species was higher in the past. For example, *V. candicans* and *V. riparia* had little predicted overlap in the present and in the Pleistocene, but substantial predicted overlap in the Holocene (**Figure 2D**). These SDMs support current or historical sympatry of species with evidence for introgression and also suggest that introgression was unlikely to have been recent.

### Introgression at chromosomal scales

For each of the nine trios with significant *D* values, we identified putative introgressed chromosomal regions by calculating *f*_d_ (Martin et al., 2015) and *f*_dM_ (M. Malinsky et al., 2015) in non-overlapping windows of 1000 SNPs along chromosomes (**Figure S7-S9**). The two metrics were significantly correlated (*R* > 0.75, *p* < 2.2e-16) along the genome (**Figure S10**) and gave qualitatively similar results; we focus on *f*dM because positive values are directionally interpretable as exchange from donor to receptor (M. Malinsky et al., 2015). For each trio we defined putative introgressed regions (pIRs) within receptor species (**Dataset S1**) as the top *f*dM windows that summed to the genomic proportion estimated by the *f*4-ratio (**Table 1 & Figure 2C**).

With pIRs identified, we evaluated their basic characteristics. For example, the mean length of pIRs within receptor genomes ranged from 154 Kb in the AGR trio (see **Table 1** for trio definitions) to 271 kb in GRM, with the number of pIRs ranging from 54 to 269 (**Table 1**). pIRs contained from 455 to 2,282 genes, depending on the donor-receptor pair, but for 5 of 9 trios the density of genes per kb within pIRs was significantly lower than the genome-wide average (**Table 1, Figure S11 & Table S7**). We evaluated GO enrichment and found 15 terms significantly enriched in pIRs (hypergeometric test, *p*-value < 0.048, **Table S8**), including cellcell signalling (GO:0007267), signaling receptor activity (GO:0038023) and nucleotide binding (GO:0000166) terms. Finally, we assessed two additional features of pIRs. First, some studies have suggested that introgressed regions should be in genomic regions of high recombination (Schumer et al., 2018). We used a genetic map to measure recombination in cM/kb (Zou et al., 2020); in every trio, pIRs are in regions of higher recombination relative to the genomic average (**Table 1, Table S9**). Second, we sought to determine if pIRs overlap with inferred selective sweeps in the recipient species (see Methods), focusing on 10 kp windows within the highest 5% of Composite Likelihood Ratio (CLR) support. On average 12.2% of pIRs had at least one putative sweep across the trios. However, sweeps were generally not enriched within pIRs; only one trio (ARC) had significantly more sweeps in pIRs than expected at random (t-test, p < 2.2e-16, **Table 1**). This trio provided an interesting example on chromosome 16, where the pIR at 14-15 Mb contained multiple putative sweeps (**Figure 3**).

### pIRs are enriched for resistance functions

pIRs were not enriched for selective sweeps, but we also examined the potential relationship between pIRs and biotic resistance. To do so, we subjected all 28,259 predicted genes to a pipeline designed to identify disease resistance genes (Osuna-Cruz et al., 2018). We tallied genes in the four most-studied types of pathogen recognition genes (PRGs) - i.e., CC-NB-LRR (CNL), TIR-NB-LRR (TNL), Receptor Like Proteins (RLP), and Receptor Like Kinases (RLK) genes. We found 208 CNLs, 55 TNLs, 258 RLKs, and 336 RLPs in the *V. arizonica* reference (**Dataset S2**), representing 3% of all genes. We then assessed whether pIRs were enriched in PRGs by permuting functions among genes. Notably, PRGs were enriched within the pIRs of receptor species for seven trios, with five significantly enriched (hypergeometric test, *p*-value < 3.66e-04) (**Figure 4A, Table 1**). Regions introgressed from *V. candicans* into *V. riparia* were especially noteworthy, because all four PRG types were significantly enriched. Illustrative examples include the pIRs at 3 Mb and 22 Mb on chromosome nine, which had 53.7% and 33.3% of their genes annotated as PRGs, respectively (**Figure 4B**). The pIR at 14 Mb in chromosome 16 of the ARC trio was another region of interest (**Figure 3**), because it contained multiple selective sweeps and also had 39.1% of the genes annotated as PRGs.

**Figure 4.**
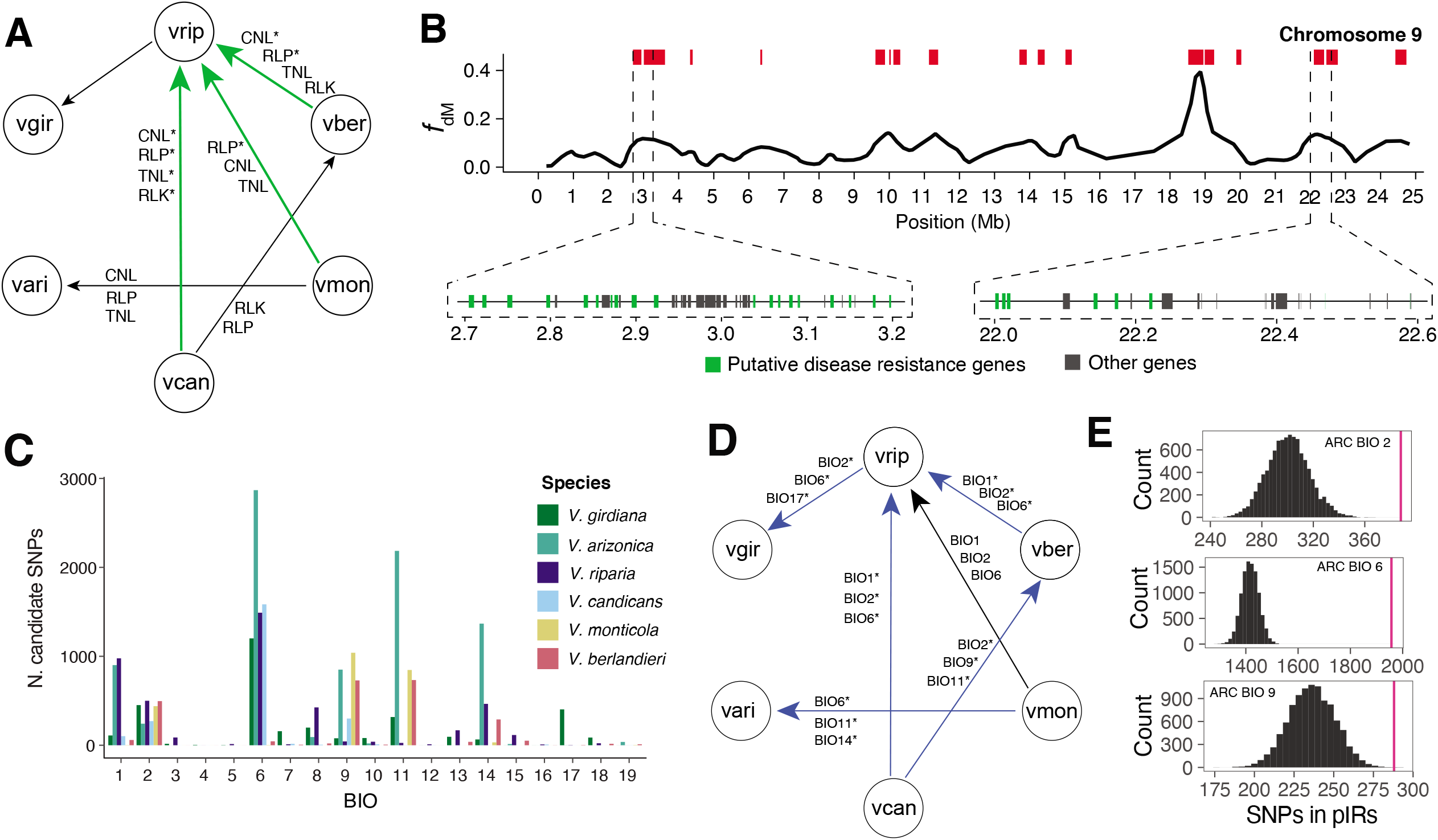
Biotic and abiotic signals in pIRs. (A) Illustrates the pIRs from donor to receptor that were significantly enriched (green) in at least one category of disease resistance genes. The four categories of disease resistance genes are CNL: CC-NB-LRR, TNL: TIR-NB-LRR, RLP: Receptor Like Proteins, and RLK: Receptor Like Kinases. (B) Shows two pIRs from V. candicans into V. riparia (ARC) in chromosome 9 with selective sweeps and highly enriched in disease resistance genes (in green). (C) Number of SNPs associated with all 19 bioclimatic variables per species. (D) Shows the pIRs from donor to receptor that were significantly enriched (blue) in at least one of the top three bioclimatic variables. (E) Diagram showing the distribution resulting of the permutations (black) in comparison with the observed value (magenta) of the top three bioclimatic variables for the ARC trio. Variables in A and D with an asterisk indicate a significant enrichment in the pIRs for the specific introgression.

The enrichment for PRGs suggests that pIRs could provide an adaptive benefit to biotic challenges. To pursue this idea further, we gathered PD resistance data for 79 accessions from the four receptor species (Krivanek et al., 2006; S. Riaz et al., 2018) (See Methods) (**Figure 1**), using variation in bacterial levels as a quantitative phenotype (**Table S10**). Bacterial levels were significantly associated (-log10(*p*) > 8) with from as few as 356 SNPs in *V. girdiana* to as many as 5,424 SNPs in *V. berlanderi* (**Dataset S3**). Most of these SNPs (64% on average, across the four species) were outside coding regions (**Table S11**). However, numerous genes contained associated SNPs, ranging from 43 in *V. girdiana* to 842 in *V. berlandieri.* Interestingly, 19 genes had associated SNPs in at least two receptor species (**Table S12**). These 19 genes, including two disease defense genes, constitute a list of candidate genes that moderate to bacterial levels. We also used these data to assess whether pIRs were enriched for associated SNPs. Depending on the trio, we detected from 0 to 323 associated SNPs in the pIRs of receptor species, and pIRs were significantly enriched for associated SNPs in four out of the nine trios (*p* < 2.2e^-16^) (**Table 1, Table S13 & S14**). This pattern occurs despite the fact that pIRs tend to be gene-poor regions of the genome.

### pIRs are enriched for environmental adaptations

pIRs may also play a role in abiotic adaptation. To evaluate this hypothesis, we examined correlations between SNPs and 19 bioclimatic variables based on Baypass analyses (Gautier, 2015), defining candidate SNPs as those with a BayesFactor (BF) >10 (Gautier, 2015; Jeffreys, 1998). The number of candidate SNPs associated with climate varied among species, ranging from 2,300 in *V. candicans* to 8,576 in *V. arizonica* (**Dataset S4**). For each species, we identified the three bioclimatic variables that had the strongest BF values (**Figure 4C, Figure S12**). Most species had candidate SNPs associated with low temperatures during the coldest period (BIO11 and BIO6), the temperature of the driest months (BIO9) and/or mean diurnal range (BIO2). Given candidate SNPs and three associated bioclimatic variables, we assessed whether pIRs in receptor species were enriched for candidate SNPs. Candidate SNPs were significantly enriched in the pIRs of seven of the nine trios (*t*-test, p < 2.2e-16; **Figure 4E, Table S15**).

## DISCUSSION

We have used population genomic approaches, specifically variants of the D statistic, to characterize putatively introgressed regions (pIRs) across six wild *Vitis* species. *Vitis* is an interesting system for studying introgression because it is an example of an adaptive radiation (Z.-Y. Ma, Wen, Ickert-Bond, et al., 2018), because several species grow in sympatry (**Figures 1, 2D & Figure S4**) and because all wild species have the potential to be (or already are) agronomically important. Moreover, all *Vitis* spp. are inter-fertile, and hybrid individuals can be found in nature (Heinitz et al., 2019). To facilitate our study of historical introgression, we first generated a reference genome assembly for *V. arizonica.* One ongoing challenge to the study of *Vitis* genomes (and that of many other plant species) is their high heterozygosity and hemizygosity (Vondras et al., 2019; Zhou et al., 2019), which complicates assembly (Minio, Massonnet, Figueroa-Balderas, Castro, et al., 2019). To date, the only wild grape genome that has been sequenced is from *V. riparia* (Girollet et al., 2019). Our *V. arizonica* genome is more complete and contiguous (e.g., scaffold N50 of 1Mb vs. 25.9Mb), presenting a clear advantage for its use as a reference.

Given our dataset of ~20 million SNPs across six species, the first goal was to infer phylogenetic history. Given the rapid radiation and inter-fertility of species in the genus, it is not surprising that phylogenetic treatments of *Vitis* have not yet reached a broad consensus (e.g. (Miller et al., 2013)). With our whole-genome dataset we have been able to recognize and remove 19 individuals of recent hybrid origin, to identify distinct monophyletic clades for each species and to produce a well-supported species-level phylogeny (**Figure 2**). Although phylogenetic inference has not been our primary focus, these results suggest that additional within-species sampling and on whole genome resequencing will provide further resolution to phylogenetic relationships within this critical genus.

Our phylogenetic analyses also indicate that individual genomic segments often do not conform to the consensus phylogeny (**Figure 2B**), suggesting the possibility of an extensive history of introgression. However, the genomic extent of introgression has not been clear for most taxa, because surprisingly few studies of introgression in plants or animals have been based on multiple species, multiple individuals per species and full genome resequencing (Payseur & Rieseberg, 2016). Here we have applied *D* and *f* statistics to test for signals of introgression against the null hypothesis of lineage sorting. Our results show that signals of introgression are common, because we detect signals between six distinct pairs of donor-recipient species among eight tested pairs (**Figure 2**). These analyses suggest that from 2% to 8% of recipient genomes owe their origins to introgression (**Figure 2 & Table 1**). The higher value, of 8%, is similar to the ~8 to 10% of the human genome inferred to have come from neanderthals and exceeds the percentage of genome (5%) genome introgressed between *Z. mays* ssp. *mays* and ssp. *mexicana* (Gonzalez-Segovia et al., 2019). Our work extends the growing recognition that a history of introgression has shaped an appreciable proportion of extant genomes.

It remains difficult, however, to infer the evolutionary forces that lead to the retention of introgressed regions within hybrid individuals (Barton & Hewitt, 1985). One hypothesis is that they are retained because they reduce genetic load, particularly if deleterious alleles are recessive. Under this model, the introgressed region has lower load in the hybrid compared to the recipient species, creating a fitness advantage. This scenario may be most likely for regions of the genome that: *i*) are unlikely to contribute numerous deleterious mutations (and hence likely to be gene poor); *ii*) have high recombination rates, where interference among mutations is minimized (Schumer et al., 2018); and *iii*) come from donor populations with higher effective population sizes (*N--e*) than recipient species. This last point reflects the fact that high *N--e* species are generally expected to have a lower deleterious load. In our data, we find no obvious evidence to suggest that *N--e* (as approximated by genome-wide *π*) is consistently higher for donor than receptor species (**Table S5**), but our results are consistent with the first two points. We find, for example, that pIRs are often gene poor, because pIRs in 5 of 9 trios had significantly lower gene density than randomly chosen genomic regions of the same size (**Table 1**). Here we must recognize an important caveat: our trios are neither evolutionary nor statistically independent. As a consequence, making conclusions based solo on the number of trios may be misleading. Nonetheless, some trends are clear; none of the nine trios have any evidence for enriched gene density within pIRs, and the pIRs in all trios are found in regions with higher recombination rates than the genome-wide average (**Table 1**).

Recent work also suggests that a history of adaptation may be necessary to produce positive introgression statistics, particularly for regions like our pIRs that have high recombination rates and low gene density (Zhang et al., 2020). To investigate potential signals and causes of adaptation in our data, we have performed two distinct types of analyses. The first is selective sweep mapping within recipient species. While there were some compelling examples of overlapping sweeps and pIRs (**Figure 3**), we do not find that pIRs are enriched for selective sweeps (**Table 1**). The lack of enrichment does not completely nullify an adaptive explanation for introgression, because selective sweep mapping is inherently noisy and because sweeps are detectable only over a finite time frame. We believe, in fact, that most of our pIRs reflect historical events, based on two sources of information. First, most species pairs that show evidence for introgression do not currently grow in sympatry; SDMs suggest they were sympatric only in the past (**Figure 2D & Figure S4**). Second, some pIRs are shared among trios. If we consider, for example, two trios that have *V. riparia* as the receptor with two different donors (*V. candicans* and *V. berlandieri*), we find 48 overlap between trios, suggesting they predate the speciation of *V. candicans* and *V. berlandieri*.

Another approach to assess adaptation is to investigate potential function. We have assessed gene content and explored associations with biotic and abiotic variables. For gene content, the pIRs have a consistent over-enrichment of genes involved in defense functions; 7 of 9 trios have a receptor species with pIRs that have more than the expected number of disease resistance genes, with 5 of 7 statistically significant (**Table 1**). This phenomenon is most pronounced for *V. riparia*, because it is the recipient species in all five of these trios. Nonetheless, when we extend our work to include a quantitative biotic phenotype – i.e., an assay of *X. fastidiosa* quantity after infection - we find that 4 of 9 trios have pIRs enriched for SNPs associated with bacterial level. Interestingly, three of these four do not include *V. riparia* as the recipient species, so that our joint approach based on gene content and PD association implicates pIRs in disease function from eight of the nine trios (**Table 1**). Altogether, based on these results, we hypothesize that pathogen resistance is a major feature that shapes the adaptive retention of introgressed regions. These results complement findings that plant-pathogen interactions play dominant roles in shaping the genetic diversity (Laine et al., 2011; Marden et al., 2017; Stump et al., 2020) and evolutionary dynamics (Bever et al., 2015; Burdon et al., 2006; Stump et al., 2020) of plant populations.

Because PD is of substantial and agronomic and economic interest, the bacterial associations merit additional discussion. First, it is worth noting that our sample sizes are small, and hence once must be cognizant of the potential for false positives. We have no reason to believe, however, that false positives should be enriched in pIRs. Second, we note that we found few associations in the region of chromosome 14 where *PdR1* is likely to be located. This region requires further study, but we believe that the *V. arizonica* genome reference does not contain a functional *PdR1* gene. Recent studies have suggested that other loci contribute to PD resistance and that these loci may vary among species (S. Riaz et al., 2018; Summaira Riaz et al., 2020). Consistent with that observation, we have found SNPs associated with bacterial levels across the genome. The number of associated genes varies widely across the four receptor species. In *V. berlandieri*, for example, we have detected associated SNPs in 842 genes, but only 43 genes in *V. girdiana*. While this difference undoubtedly reflects that statistical power differs among species, there are some consistent signals, such as the 19 genes that are SNP associations in more than one species. There are a few compelling candidates among the 19 genes, including an NBS-LRR receptor kinase gene (*g270290*) on chromosome 19, and a gene containing an NB-ARC domain (*g167550*) in chromosome 12 (**Table S12**). We propose that these genes contribute in some way to *X. fastidiosa* resistance or tolerance, but they require future functional study.

We have also assessed the potential for adaptive introgression by associating genetic diversity with bioclimatic variables; climate-associated SNPs were highly significantly enriched within pIRs (p < 2.2 e-16) for seven of the nine trios. A common theme is that cold-adapted alleles appear to have introgressed into *V. riparia*. This is a slightly puzzling result, both because *V. riparia* is the most cold hardy species among North America wild *Vitis* (Pierquet & Stushnoff, 1980) and also because our sample represents a geographic region that is likely to experience warmer temperatures than most *V. riparia*. However, recent studies about the physiological processes involved in grape cold hardiness may provide clues to this puzzle (Kovaleski et al., 2018; Londo & Kovaleski, 2019). Perennial plants reduce cold hardiness in a process known as deacclimation that results in budbreak is critical for their survival, and deacclimation rates vary among wild grape species (Kovaleski et al., 2018). It is possible that historical introgressions into *V. riparia* facilitated adaptation to the southern regions by receiving alleles involved in different mechanisms of cold adaptation – e.g., involved in the avoidance of premature bud-break during midwinter increases of temperature. There is another intriguing possibility: given that pIRs are biased for resistance genes, especially in *V. riparia*, climate-associations may not reflect adaptation to climate directly but rather indirect indicators of biotic stresses within a particular climate. Some evidence from this speculation comes from our observation that, on average, 17.8% of the genes annotated as PRGs in pIRs had a candidate SNP associated with a bioclimatic variable.

Altogether, our work contributes to the idea that introgression contributes phenotypic innovations and evolutionary novelty (Arnold & Kunte, 2017) by reassorting genetic variants into beneficial combinations that permit adaptation to new ecological niches (Marques et al., 2019). In this specific case, in *Vitis* from the Southwestern United States, it suggests that adaptive introgression has been driven by responses to biotic stress (disease) and perhaps abiotic stress (climate). On a practical scale, our inferences may help identify genetic variants that will prove to be valuable for rootstock breeding.

## MATERIALS and METHODS

### *V. arizonica* genome

#### DNA library preparation and sequencing

High-molecular-weight genomic DNA (gDNA) was isolated using the method described in Ref. (Chin et al., 2016). DNA purity was evaluated with a Nanodrop 2000 spectrophotometer (Thermo Scientific, Hanover Park, IL, USA), DNA quantity with the DNA High Sensitivity kit on a Qubit 2.0 Fluorometer (Life Technologies, Carlsbad, CA, USA), and DNA integrity by pulsed-field gel electrophoresis. gDNA was cleaned with 0.45x AMPure PB beads (Pacific Biosciences, Menlo Park, CA, USA) before library preparation. SMRTbell template was prepared with 15 μg of sheared DNA using SMRTbell Template Prep Kit (Pacific Biosciences, Menlo Park, CA, USA) following the manufacturer’s instructions. SMRTbell template was size-selected using the Blue Pippin instrument (Sage Science, Beverly, MA, USA) using a cutoff size of 17-50 Kbp. The size-selected library was cleaned with 1x AMPure PB beads followed by a DNA damage repair treatment and cleaned again with 1x AMPure PB beads. The three SMRTbell libraries produced were sequenced in 12 SMRT cells on a PacBio Sequel I platform (DNA Technology Core Facility, University of California, Davis).

#### Genome assembly

The *V. arizonica* genome was assembled using a hybrid strategy combining Single Molecule Real-Time (SMRT; Pacific Biosciences) with Bionano NGM maps (Bionano genomics). Genomic sequences were assembled using SMRT reads following the custom procedure reported in https://github.com/andreaminio/FalconUnzip-DClab (Minio, Massonnet, Figueroa-Balderas, Castro, et al., 2019). The pipeline performs the marking of repetitive content in SMRT reads using the TANmask and REPmask modules from DAmasker suite (Myers, 2014), both on raw reads before error correction and on the corrected reads used by FALCON ver. 2017.06.28-18.01 (Chin et al., 2016) to assemble the genome sequences. Multiple combinations of assembly parameters were tested to reduce the sequence fragmentation. We used the following parameters for the final assembly:

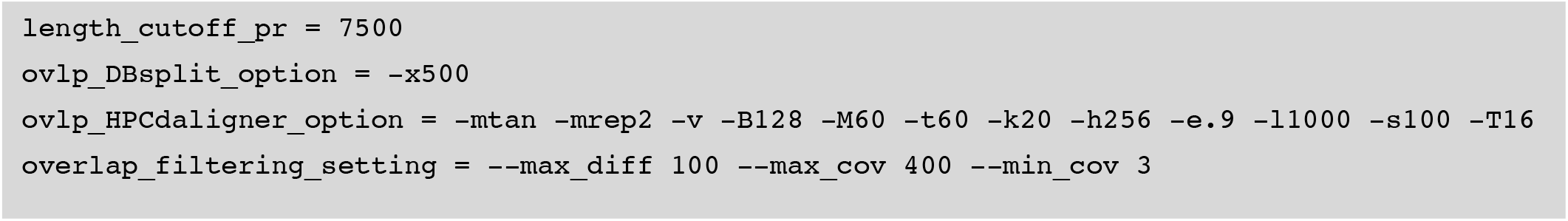

Unzip with default parameters was performed to produce a diploid representation of the genome with primary contigs and the associated phased haplotigs (Chin et al., 2016) that were then polished from sequence error using PacBio reads with Arrow (from ConsensusCore2 ver. 3.0.0). Primary assembly contiguity was improved by scaffolding with SSPACE-Longread ver. 1.1 (Boetzer & Pirovano, 2014), followed by a gap-closing procedure with PBJelly (PBsuite ver. 15.8.4) (English et al., 2012, 2014). Bionano molecules (Bionano Genomics) were generated and assembled at the genome center of the University of California, Davis (Ming-Cheng Luo). Labeling was performed with DLE-1 non-nicking enzyme (CTTAAG) and molecules were then sequencing on a Saphyr machine. Optical maps were assembled with BioNano Solve ver. 3.3 (Lam et al., 2012). The optical maps obtained were then used to scaffold the PacBio assembled sequences using HybridScaffold ver. 04122018 (Lam et al., 2012). The procedure was performed in four iterations. In the first iteration both sequences and optical maps were broken (‘-B 2 -N 2’) when in conflict one with the other. In the second iteration, the scaffolds produced as results of the first iteration were compared to the optical maps, again both sequences and optical maps were broken (‘-B 2 -N 2’) when in conflict. In the third iteration, scaffolds resulting from the previous step were compared to the optical maps, conflicts were resolved by breaking nucleotide sequences (‘-B 1 -N 2’). In the fourth iteration, results of the previous scaffolding were used, conflicts were again resolved by breaking nucleotide sequences (‘-B 1 -N 2’). The genomic sequences obtained were organized and sorted into two sets of chromosomes by using HaploSync ver. 0.1beta (https://github.com/andreaminio/HaploSync) and based on synteny with *Vitis vinifera* PN40024 chromosomes. The genome is available at http://www.grapegenomics.com/pages/Vari/

#### Genome annotation

The structural annotation of the *V. arizonica* genome was performed with the pipeline described here: https://github.com/andreaminio/AnnotationPipeline-EVM_based-DClab, an adaptation of the one presented in Ref. (Vondras et al., 2019) to make use of IsoSeq data as primary experimental evidence. In brief, high-quality Iso-Seq data from *V. arizonica* were used in PASA ver. 2.3.3 (B. J. Haas, 2003) to generate a set of high-quality gene models for the training of the following ab initio predictions software: Augustus ver. 3.0.3 (Stanke et al., 2006), GeneMark ver. 3.47 (Lomsadze, 2005) and SNAP ver. 2006-07-28 (Korf, 2004). Gene predictions were generated also using BUSCO ver. 3.0.2 (Simão et al., 2015) with OrthoDB ver. 9 Plant conserved proteins. Repeats were annotated using RepeatMasker ver. open-4.0.6 (Smit, AFA et al., 2015) with the *Vitis* custom repeat library reported in Ref. (Minio, Massonnet, Figueroa-Balderas, Vondras, et al., 2019).

Transcriptomic data was obtained from Ref. (Minio, Massonnet, Figueroa-Balderas, Vondras, et al., 2019), Vitis ESTs (NCBI, download date: 2016.03.15) and Vitis mRNA excluding transposable elements related proteins (NCBI, download date: 2016.03.15). RNA-Seq was processed using Stringtie ver. 1.3.4d (Pertea et al., 2015) and Trinity ver. 2.6.5 (Grabherr et al., 2011) with both on-genome and de novo protocols. Transcriptomic data was mapped, along with transcriptome assemblies and from the Iso-Seq data described above, on the *V. arizonica* genome using PASA ver. 2.3.3 (B. J. Haas, 2003) and MagicBLAST v.1.4.0 (Boratyn et al., 2019). Protein evidence obtained from swissProt viridiplantae (download date: 2016.03.15) and Vitis proteins excluding transposable elements related proteins (NCBI, download date: 2016.03.15) were mapped on the genomic sequences using Exonerate ver. 2.2.0 (Slater & Birney, 2005). Predictions and experimental evidence were then processed by EvidenceModeler ver. 1.1.1 (Brian J Haas et al., 2008) to generate consensus gene models and alternative splicing information integrated from the available transcriptomic data using PASA ver. 2.3.3 (B. J. Haas, 2003). The final functional annotation was produced integrating into Blast2GO ver. 4.1.9 (Conesa et al., 2005) hits from the blastp ver. 2.2.28 (Altschul et al., 1990) results against the Refseq plant protein database (ftp://ftp.ncbi.nlm.nih.gov/refseq, retrieved January 17th, 2017) and InterProScan ver. 5.28-67.0 (Jones et al., 2014).

Disease-related gene functions were annotated using the HMM models from the Disease Resistance Analysis and Gene Orthology (DRAGO 2) database (Osuna-Cruz et al., 2018). All the 28,259 predicted proteins from haplotype 1 anchored in the 19 chromosomes were evaluated in DRAGO2. Enrichment analysis of functional categories was tested using GeneMerge v1.4 (Castillo-Davis & Hartl, 2003).

#### cDNA library preparation and sequencing

Total RNA from *V. arizonica* leaves was isolated using a Cetyltrimethyl Ammonium Bromide (CTAB)-based extraction protocol as described in Ref. (Blanco-Ulate et al., 2013). Nanodrop 2000 spectrophotometer (Thermo Scientific, Hanover Park, IL) was then used to evaluate RNA purity. The RNA quantity was evaluated with the RNA broad range kit of the Qubit 2.0 Fluorometer (Life Technologies, Carlsbad, CA) and the integrity using electrophoresis and an Agilent 2100 Bioanalyzer (Agilent Technologies, CA). Total RNA (300 ng, RNA Integrity Number > 8.0) was used for cDNA synthesis and library construction. An RNA-Seq library was prepared using the Illumina TruSeq RNA sample preparation kit v.2 (Illumina, CA, USA) following Illumina’s Low-throughput protocol. This library was evaluated for quantity and quality with the High Sensitivity chip in an Agilent 2100 Bioanalyzer (Agilent Technologies, CA) and was sequenced in 100bp single-end reads, using an Illumina HiSeq4000 sequencer (DNA Technology Core Facility, University of California, Davis). To prepare a cDNA SMRTbell library, first-strand synthesis and cDNA amplification were accomplished using the NEBNext Single Cell/Low Input cDNA Synthesis & Amplification Module (New England, Ipswich, MA, US). The obtained cDNAs were then purified with ProNex magnetic beads (Promega, WI) following the instructions in the Iso-Seq Express Template Preparation for Sequel and Sequel II Systems protocol (Pacific Biosciences, Menlo Park, CA). Amplified cDNA was size-selected with a mode of 2 kb using ProNex magnetic beads (86 μl). At least 80 ng of the size-selected, amplified cDNA was used to prepare the cDNA SMRTbell library using the SMRTbell Express Template Prep Kit 2.0 (Pacific Biosciences, Menlo Park, CA), following the manufacturer’s protocol. One SMRT cell was sequenced on the PacBio Sequel I platform (DNA Technology Core Facility, University of California, Davis).

### Genetic Diversity Data and Analyses

#### Plant material

We collected fresh leaf tissue from 130 individuals from six American *Vitis* species in the wild grape germplasm collection at Davis, California. The germplasm reported in this study was collected from 1997 to 2016 as cuttings across the southwestern states and maintained at the Department of Viticulture and Enology, University of California, Davis, CA. Table S1 provides details of global positioning coordinates and species designation based on the morphological features of leaves and growth habit of the field grown plants.

#### Genome resequencing

For the 130 samples, genomic DNA was extracted from leaf samples with the Qiagen DNeasy plant kit. The sequencing libraries were constructed with an insert size of ~300 bp using Illumina library preparation kits and were sequenced using the Illumina HiSeq 2500 platform with 2 × 150 bp paired reads to a target coverage of 25X following ref. (Zhou et al., 2017). The raw sequencing data has been deposited in the Short Read Archive at NCBI under BioProject ID: PRJNA607282.

#### SNP call and filtering

We filtered and evaluated raw reads using Trimmomatic-0.36 (Bolger et al., 2014) and FastQC (http://www.bioinformatics.babraham.ac.uk/projects/fastqc/). Filtered reads were then mapped to the reference genome with the BWA-MEM algorithm (Li, 2013) implemented in bwa-0.78 (Li, 2014). Joint SNP calling was conducted using the HaplotypeCaller in the GATK v.4.0 pipeline following ref. (Zhou et al., 2017). We then filtered raw SNPs with bcftools v1.9 (https://samtools.github.io/bcftools/) and vcftools v0.1.15 (https://vcftools.github.io/). We kept SNPs sites for downstream analysis if they were biallelic, had quality higher than 30, had a depth of coverage higher than five reads, also had no more than three times the median coverage depth, and had less than 75% of missing data. Additionally, the following expression was applied under the exclusion argument of the filter function in bcftools: “QD < 2.0 | FS > 60.0 | MQ < 40.0 | MQRankSum < −12.5 | ReadPosRankSum < −8.0 | SOR > 3.0”.

#### Population Structure and Phylogenetic Analyses

We used NgsAdmix from the ANGSD package v0.915 (Skotte et al., 2013) to evaluate genetic structure and measure genetic diversity (π) within species. We tested values of *K* from 1 to 10 and used a minMaf filter of 0.05. We then employed the Cluster Markov Packager Across K (Clumpak) software (Kopelman et al., 2015) to detect the best *K*, which resulted in *K*=7. For the overall consensus phylogenetic tree, we used the SNPhylo pipeline (v20180901) to reduce the number of SNPs (Lee et al., 2014), using a linkage disequilibrium threshold of 0.5 which resulted in 15,893 informative sites. We then created a maximum likelihood phylogeny with the sites from SNPhylo using IQ-TREE v1.6.12 (Nguyen et al., 2015). We used the Model Finder algorithm (Kalyaanamoorthy et al., 2017) implemented in IQ-TREE to search for the substitution model best among 550 different combinations. The combination of models used was “VT+F+R5”. We used the ultrafast bootstrap option with 1000 replicates to obtain support values for each node. For the densitree (in red, Figure 1C) we divided the genome in 10 Kb windows but focused on used windows with at least 800 variable sites, resulting in 3,342 windows. We created a consensus tree for each of the 3,342 windows with 1000 replicates. We then focused trees with a median bootstrap supper higher than 70% across all nodes, reducing the number of windows to 1,172, and randomly chose 500 of the 1,172 trees for plotting. We used the R packages ape v5.4 (Paradis & Schliep, 2019) and phangorn v2.5.5 (Schliep et al., 2017) to create the densitree plot and calculate tree statistics.

#### Admixture proportions

To test for introgression, we used the ABBA-BABA test as part of the Dsuite v0.4 software (Milan Malinsky et al., 2020). First, we used the program Dtrios to calculate the overall *D* statistic and perform a block jackknifing of the statistic to obtain an associated *p*-value. We used the non-hybrid samples of each species as populations. We used all *V. girdiana* samples together given their strong phylogenetic grouping (Figure 2B). The trios that were concordant with the 4-taxon topology of the test and had a *p*-value < 0.001 were then used for more detailed analysis. For the *fd* and *f*dM tests of introgression, we used the program Dinvestigate from Dsuite, choosing 1000 bp SNP windows with 100 bp steps throughout the whole genome. Finally, we defined a pIR as windows with highest *x* % of *f*dM values, where *x* was determined for each receptor species in the trio by the *f*4 estimate (Table 1).

#### Selective Sweeps

To detect selective sweeps we used RAiSD v2.8 (Alachiotis & Pavlidis, 2018) for each species separately, with default parameters. We removed the gaps of the reference genome from the analysis to avoid potential errors. We split the genome into non-overlapping 10 Kb windows created by bedtools v2.27.1 (Quinlan & Hall, 2010) and focused on the windows with top 5% of the Composite Likelihood Ratio (CLR) in the corresponding receptor population and defined those as highly supported sweeps. We first calculated the number of top 5% CLR windows from the receptor species overlapping the IRs of each trio, using bedtools with a requirement of at least 50% of the window overlapping with the IR (-f 0.5). For the enrichment analysis, we randomly chose the same number of 10 Kb windows as the number of HSSs chosen from the whole genome with 10,000 replicates. We counted the number of the random windows overlapping IRs in the same way as described before. We then compared the distribution of the observed values with the randomly generated distribution and performed a t-test to evaluate a statistical significance. For plotting, we calculated the average CLR values for a 100 Kb window using the mapping function of bedtools for clearness.

#### Recombination

To estimate the location of pIRs relative to recombination rates across the genome, we employed the genetic mapping data from (Zou et al., 2020). Briefly, the map data defines the recombination rate between two markers, based on a consensus of four different mapping populations that used both wild and cultivated *Vitis* parents. The physical location of 1662 markers was provided on the PN40024 *V. vinifera* reference. Since the V. arizonica assembly was anchored on the PN40024 *V. vinifera* reference, we were able to transform the markers and recombination values from Ref. (Zou et al., 2020). Once discrete regions of the *V. arizonica* genome had been assigned recombination values (in cM/kb), we were able to query whether pIR regions in each receptor species tended to be in regions of high recombination relative to the genome average.

### Phenotype Data, Climate Data and SNP Associations

#### Pierce’s disease assays

Evaluations for Pierce’s disease (PD) resistance were carried out using a greenhouse-based screen (Krivanek & Walker, 2005; Summaira Riaz et al., 2020). Accessions with strong and intermediate PD resistance and inoculated and un-inoculated susceptible *Vitis vinifera* cultivar Chardonnay were used in all experiments as reference control plants. Nineteen screens were carried out from 2011 to 2020, and a minimum of four biological replicates of each accession were tested. Plants were propagated, maintained and inoculated following the procedures described in Riaz et al. 2020, which also reports a subset of the data used in this study (Table S10). Disease severity was accessed 10 to 14 weeks post inoculation and ELISA was used to measure the *X. fastidiosa* levels in the stem. Statistical analysis was performed using JMP Pro14 software (Copyright 2020, SAS Institute Inc.) to determine the variability of ELISA for the reference control plants across 19 experiments. In the next step, ELISA values of wild accessions were analyzed with the inclusion of the reference plants to adjust for variation among the screens.

#### SNP Associations with PD

For each species, we used Latent Factor Fixed Models (LFFM) to evaluate the association between SNPs and PD resistance. LFMM estimates the association between the genotypic matrix and an environmental or phenotypic predictor while controlling for unobserved latent models, such as the genetic structure between populations (Frichot et al., 2013). SNPs that are significant after controlling for latent factors are considered candidate SNPs. Here we used resistance to Pierce Disease (PD) as the predictor variable, using a quantitative measure of the bacterial amount baked on an ELISA assay. Higher concentrations of the bacteria reflect lower PD resistance.

To determine the number of latent factors (LF) in each of the 6 species, we used LEA package in R to perform a principle component analysis (PCA) and a sparse nonnegative matrix factorization (SNMF) between accessions. We performed the LFMM analyses testing three different number of latent factors (LF-1 to LF+1). We used the “lfmm.pvalues” function of LEA to transform z-scores into *p*-values and performed a Bonferroni correction to control for multiple comparisons. We defined as candidate SNPs those that were identified by the three runs using different latent factors.

Because we found that IRs were hotspots of disease resistance genes, we were interested in evaluating the distribution of the PD significantly associated SNPs (-log10(p) > 8) within pIRs. We thus selected random SNPs across the genome in sets with the same number as the PD significant SNPs in the receptor species of each trio for a total of 10,000 sets. We then intersected and counted the number of randomly generated positions that fell inside IRs and compared that distribution with the observed values of SNPs in IRs. We performed a *t*-test to assess statistical significance.

#### Species distribution models

We constructed species distribution models (SDM) to identify whether species having evidence of introgression had overlapping distributions in the present, the Holocene (~6,000 ya) and the Pleistocene (~18,000 ya). For each species, we obtained occurrence records by combining coordinates data from the Global Biodiversity Information Facility (GBIF.org, 7 August 2020). Most of the six species analyzed had adequate occurrence data (>70 entries), except *V. monticola* with 20 entries. The low values for *V. monticola* is likely to lead to an overestimation of its geographic distribution. We also obtained 19 bioclimatic variables from WorldClim (R. J. Hijmans et al., 2005) at a 2.5 resolution for the present, Holocene, and Pleistocene layers. For the past layers, we obtained bioclimatic data projected with the general circulation model CCSM.

We employed Maxent V. 3.4.1 (Phillips et al., 2006) to construct and train the SDM of each species and then projected them onto the landscape for each time period. For each species, we ran 30 bootstrap replicates and used 70% and 30% of data for training and validating the models, respectively. Additionally, we evaluated the fit of the models by analyzing the area under the curve (AUC) of the receiver-operating characteristic curve (ROC). For each species, we binarized each replicate by defining as the probability of existence all areas that had a probability value in which the omission rate of the training and testing was above the 10% of prevalence, and/or that predicted the accessions we sampled. For each species, we summed all the binarized models and defined as the final distribution all geographic areas that were predicted by >60% of the bootstrap replicates.

We used SDM to predict areas of overlap between pairs of species at different time periods. To simplify the data, for each period, first, we summed the SDM of each pair of 15 species pairs retained areas that were predicted by both species. The present, Holocene and Pleistocene overlapped regions were arbitrarily set at values of 2, 5, and 10. Second we summed the three time periods to identify the areas where each pair of species were potentially sympatric at different periods. Areas with values 2, 5, and 10 correspond to areas where there have been overlaps only in the present, Holocene, or Pleistocene, respectively. Areas with values 7, 12, or 15 are areas where there have been overlaps in the present and Holocene, present and Pleistocene or Holocene and Pleistocene, respectively. Areas with value 17 are areas where there has been a continuous overlap between the pair of species. For each pair of species, we obtained the geographic area (measured by the number of overlapping pixels) that were predicted to have an overlap in the SDMs in the different periods.

#### Bioclimatic associations

Genome environment association studies identify outlier loci that correlate with the environment while controlling for demographic structure (Tiffin & Ross-Ibarra, 2014). We used Baypass version 2 (Gautier, 2015) to identify outlier loci that correlated with 19 bioclimatic variables. Baypass uses the entire set of SNPs to estimate the covariance between populations and identify populations that are genetically closer because of recent coancestry or lower genetic structure. In a second step, Baypass analyses the correlation between the allelic frequency of each SNPs (response variable) and independent environmental, phenotypic, and/or categorical variables. SNPs are considered candidates if they show a strong correlation with the independent variable after controlling for the co-ancestry between populations.

We obtained 19 bioclimatic variables from Worldclim (R. J. Hijmans et al., 2005) for the location of each accession, using the *extract* function of the raster package (R. Hijmans & van Etten, 2020) in R (R Core Team). We ran BAYPASS using all the default parameters and the 19 bioclimatic variables. Outlier SNPs were defined using Jeffreys’ rule (Jeffreys, 1998) and were considered strong candidates if they had a Baye’s Factor between 10 and 20 and decisive candidates if they had BF > 20. For each species, we identified three bioclimatic variables that showed the greatest enrichment of outlier loci and focused on decisive candidate SNPs. We further analyzed the set of outlier SNPs to see if they were enriched in pIRs. We used bedtools v2.27.1 (Quinlan & Hall, 2010) to obtain the overlap between the candidate SNPs of the top 3 bioclimatic variables of each species and the pIRs. For each trio showing evidence of introgression, we estimated whether the receptor species was enriched with SNPs associated with any of the top three candidate SNPs at the IRs. Briefly, for each trio, we first counted the number of SNPs with BF > 10 (strong candidates). Second, we counted the number of SNPs with BF > 10 that were within introgressed regions. Third, we generated 10,000 sets of random “SNPs”, each having the same number of “SNPs” as the candidate SNPs. Fourth, we obtained the number of SNPs within IRs from the random distribution for each of the 10,000 replicates. Finally, we used a *t*-test to compare the number of candidate SNPs observed in IRs and the number of SNPs in IRs from the random distribution.

## ACKNOWLEDGMENTS

The authors thank R. Gaut for generating resequencing data, R. Figueroa-Balderas and O. Nguyen (Genome Center, UC Davis) for the technical assistance and B. Forrestal for providing comments. This work was funded by the NSF grant #1741627 to B.S.G., M.A.W. and D.C., and partially supported by funds to D.C. from the E.&J. Gallo Winery and the Louis P. Martini Endowment in Viticulture.

