## Supplemental Information for "Extensive introgression among North American wild grapes (*Vitis*) fuels biotic and abiotic adaptation"

[1] Dept. Ecology and Evolutionary Biology, University of California Irvine, Irvine, CA

[2] Dept. of Viticulture and Enology, University of California, Davis, Davis, CA

### SUPPLEMENTAL FIGURES

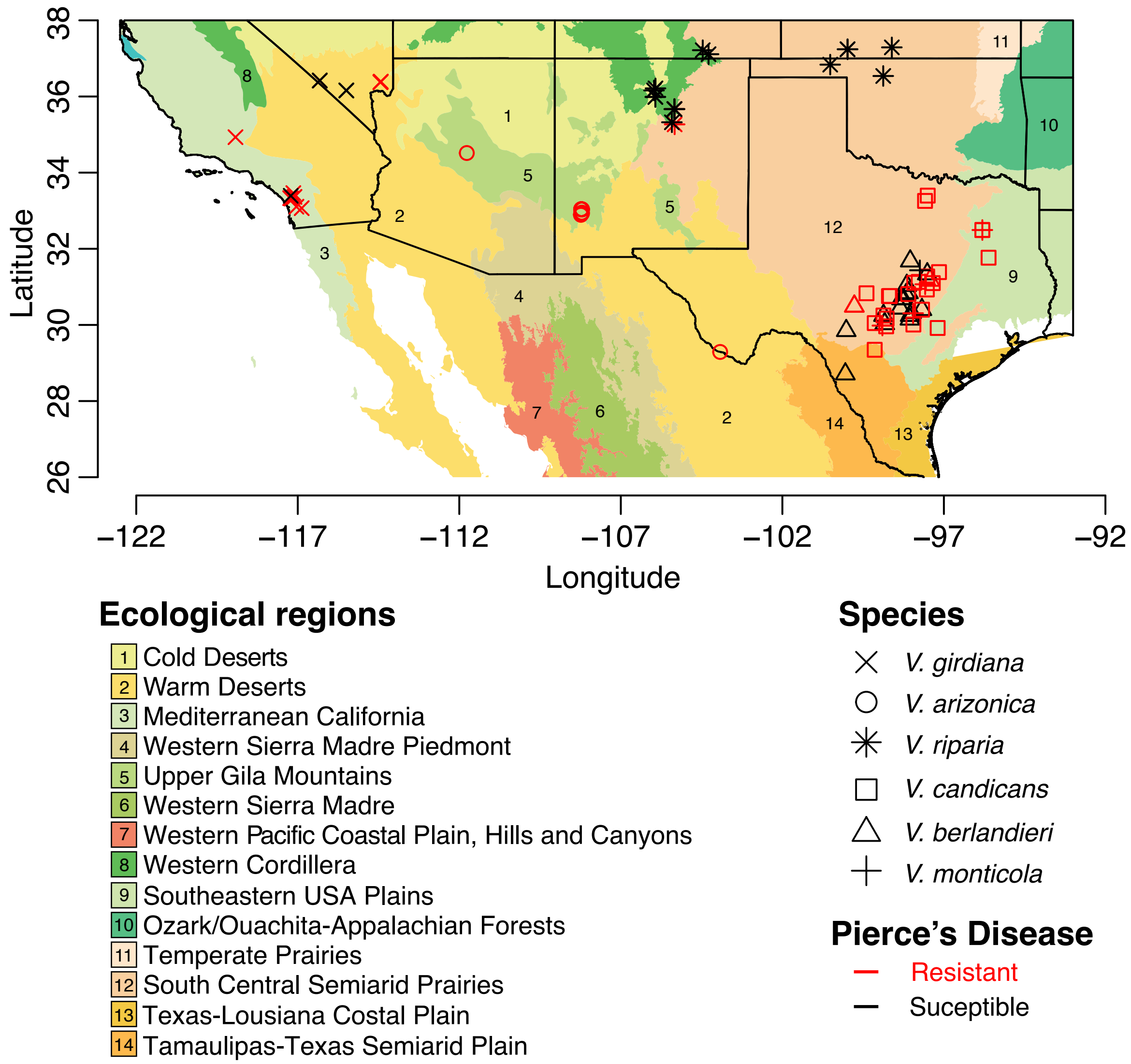


**Figure S1.** Geographic distribution and ecological diversity of sampled populations of wild grapes. Shapes correspond to different species, and samples colored red or black were classified as resistant or susceptible to Pierce's Disease, respectively. The numbers and colors of the regions correspond to the level II Ecological Regions defined by the United States Environmental Protection Agency (EPA), as shown in the legend.

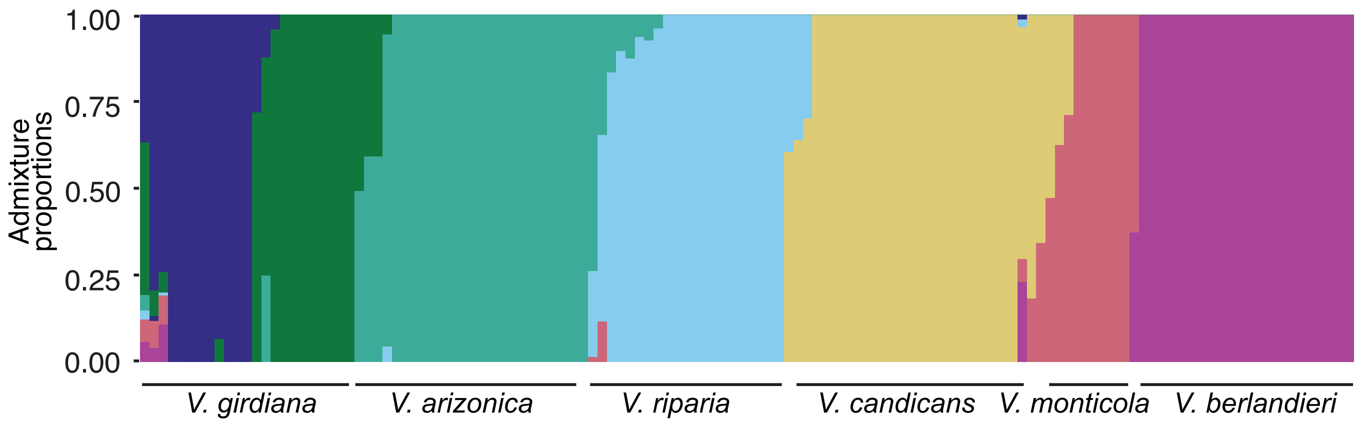


**Figure S2.** Genetic structure of all samples, including hybrid samples, with colors representing each cluster detected by the structure analysis (K=7).

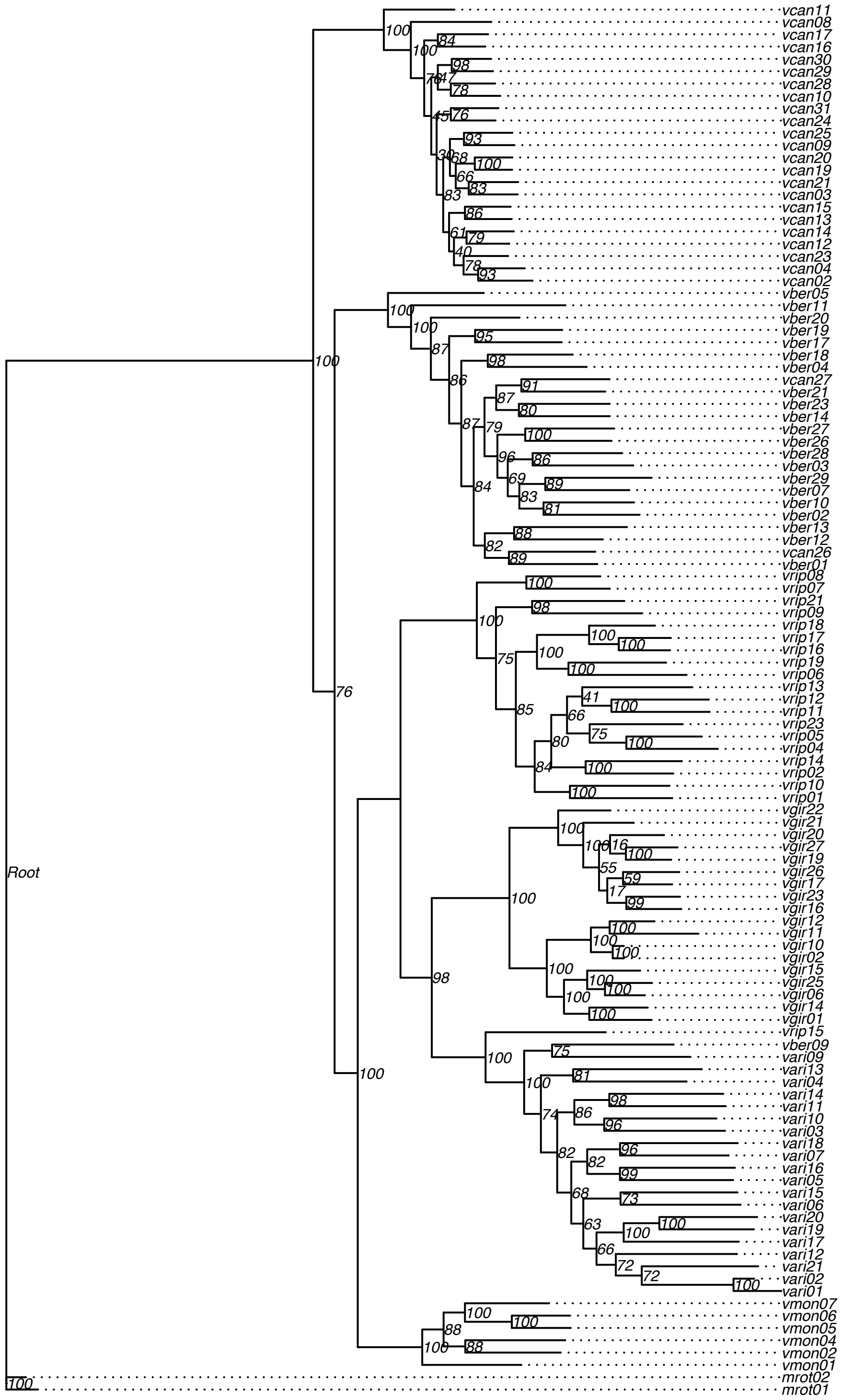


**Figure S3.** Consensus maximum likelihood phylogenetic tree created with IQtree. Numbers on the nodes correspond to bootstrap support values.

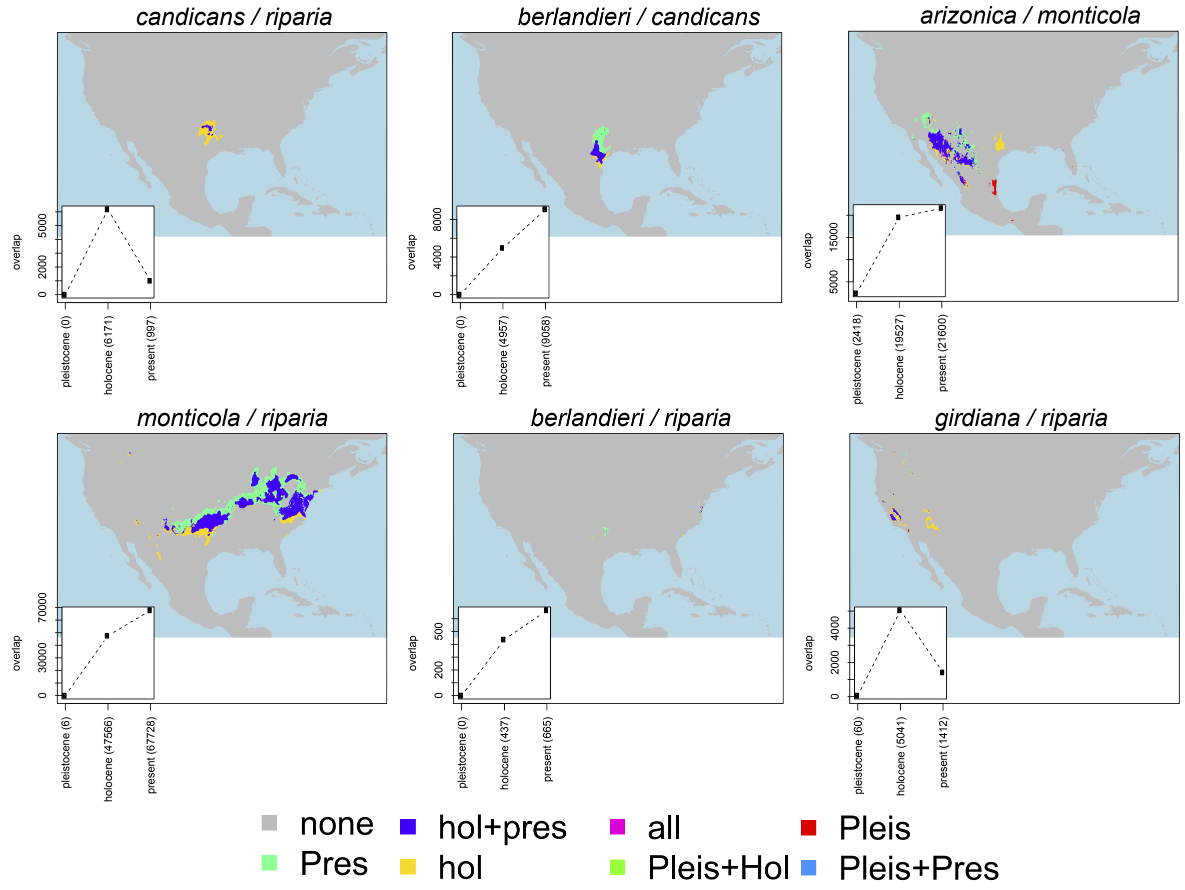


**Figure S4.** Pairs of species that show evidences of introgression present evidences of overlap at three periods, present, Holocene, Pleistocene.

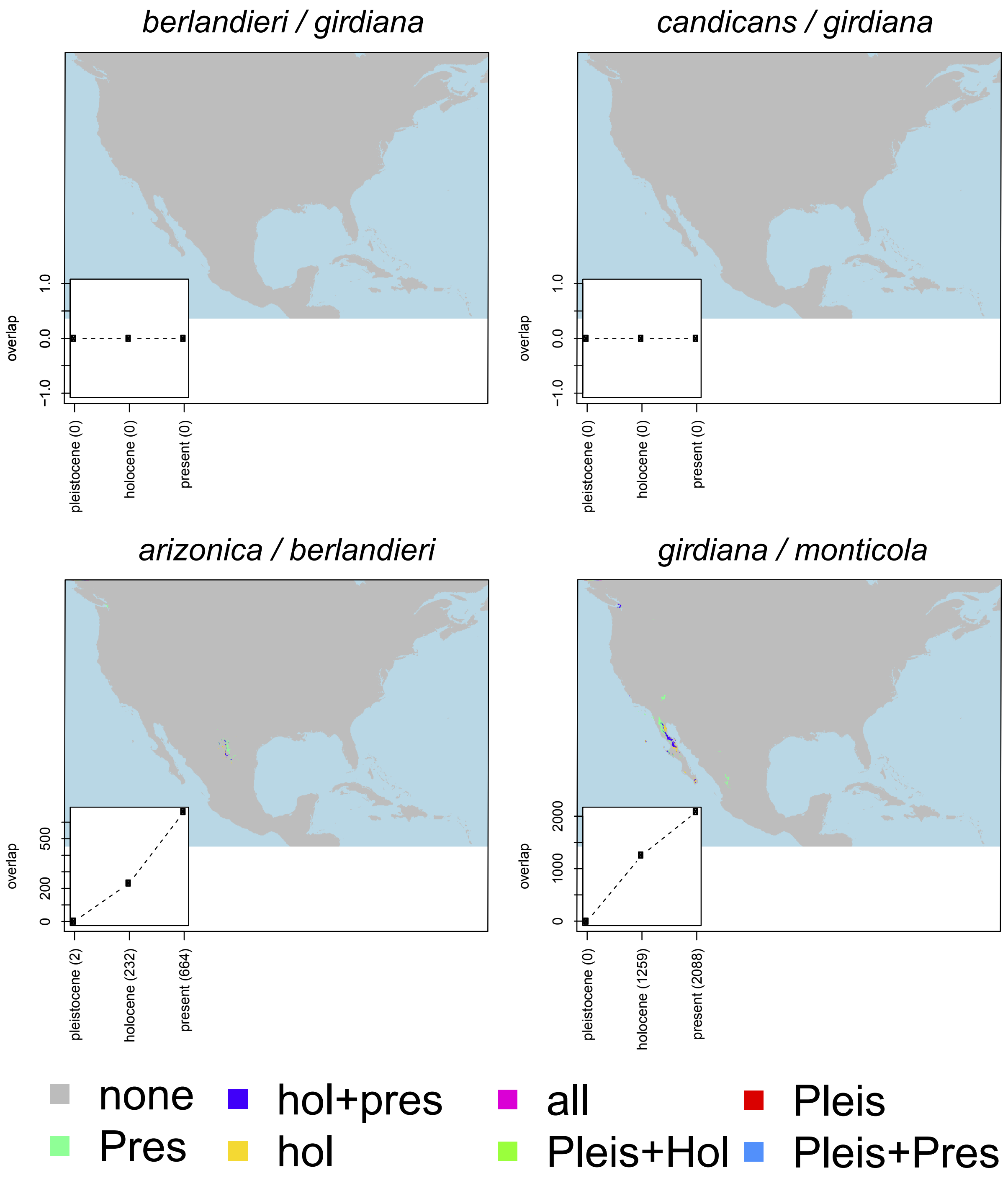


**Figure S5.** Pairs of species that do not show evidences of introgression present no or low overlap at three periods, present, Holocene, Pleistocene.

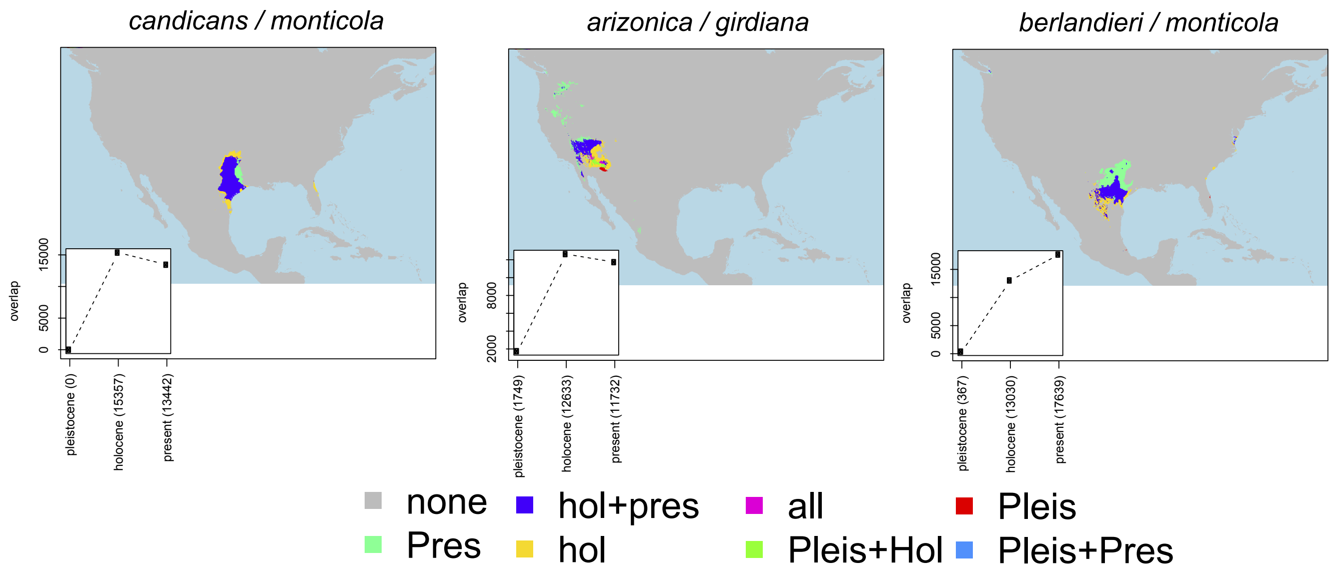


**Figure S6.** Pairs of species that do not show evidences of introgression but have overlapping distributions at three periods, present, Holocene, Pleistocene

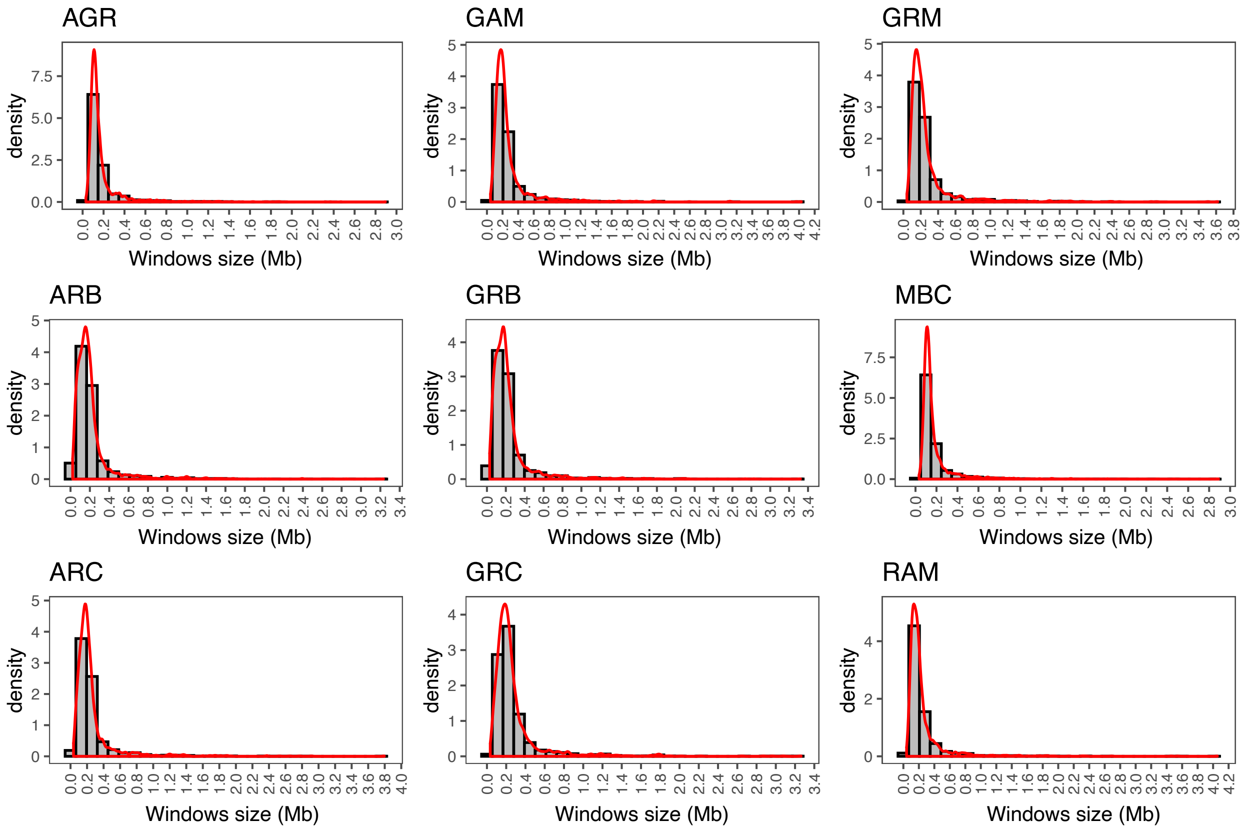


**Figure S7.** Genomic size distribution of the windows used to calculate the ABBA-BABA statistics per trio.

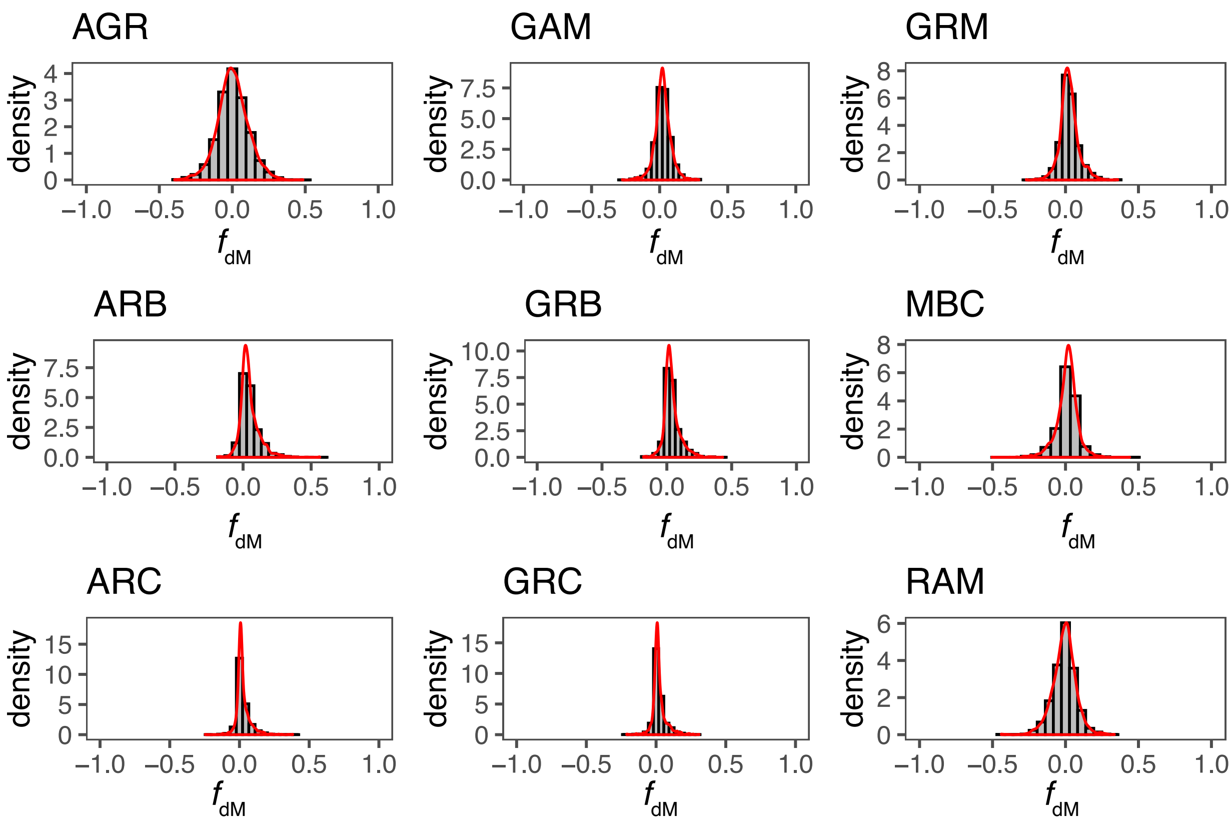


**Figure S8.** Distribution the $f_{dM}$ statistic values across windows per trio.

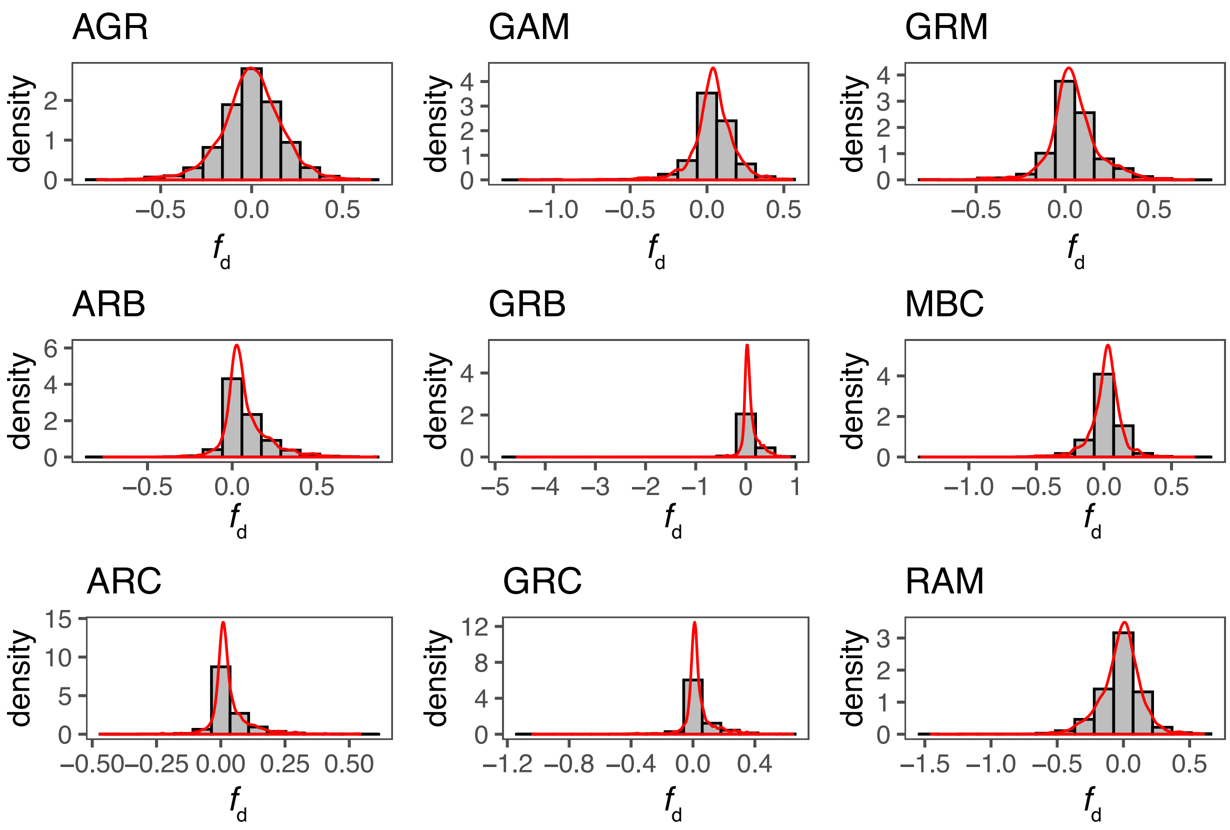


**Figure S9.** Distribution the $f_d$ statistic values across windows per trio.

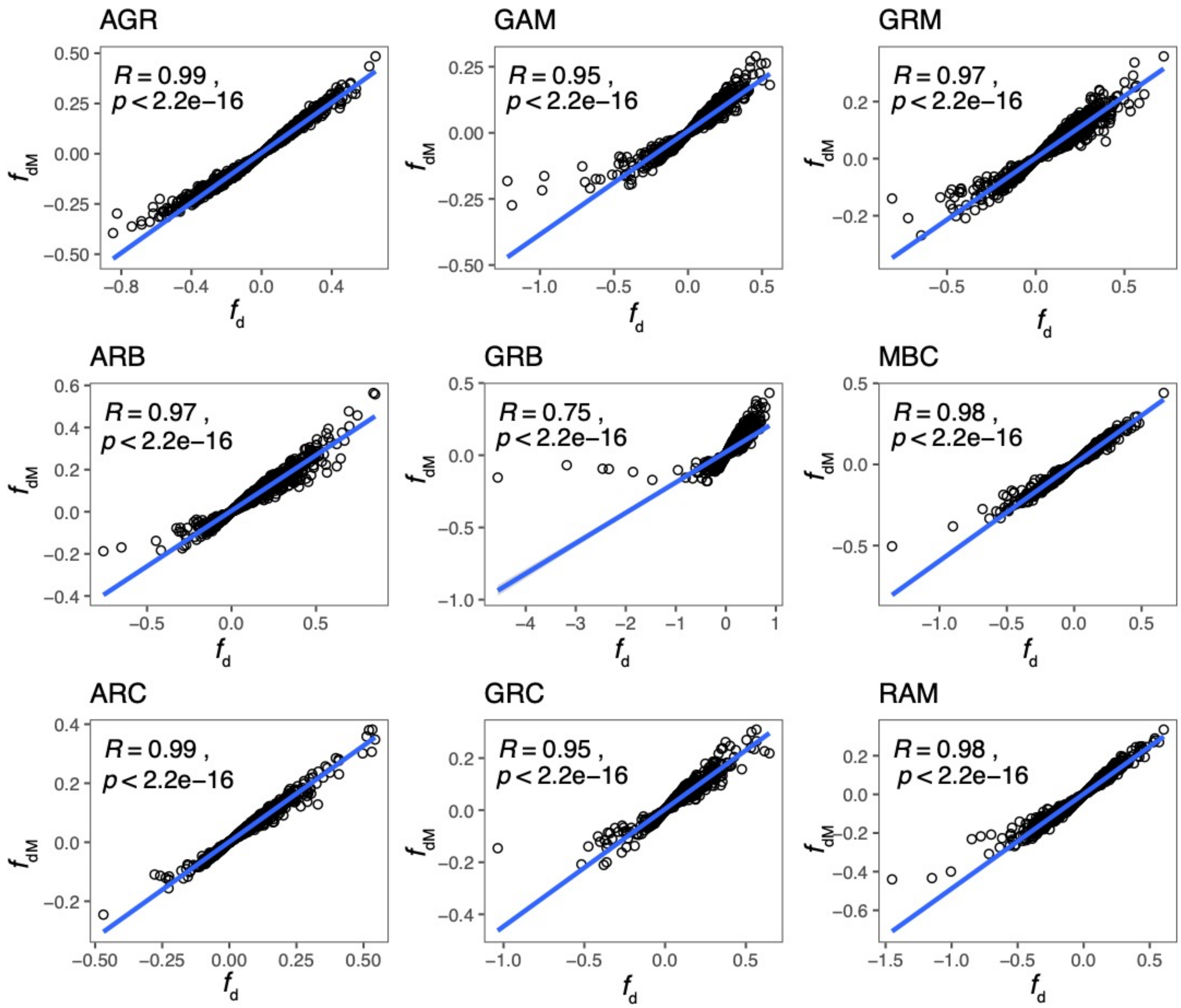


**Figure S10.** Scaterplot showing correlations between the *f*dM and *f*d statistics across all windows in the genome.

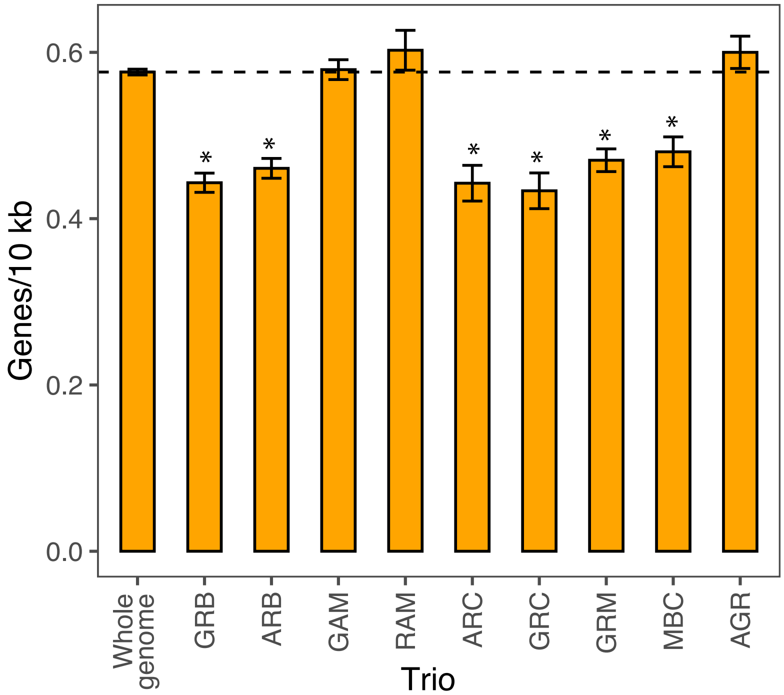


**Figure S11.** Mean gene density of across the whole genome and within the pIRs of all the trios. Error bars correspond to 1 SD. Dotted line corresponds the mean gene density of the whole genome. Asterisks denote trios with significantly lower gene density than the genome-wide average (t-test, $p$ < 0.001).

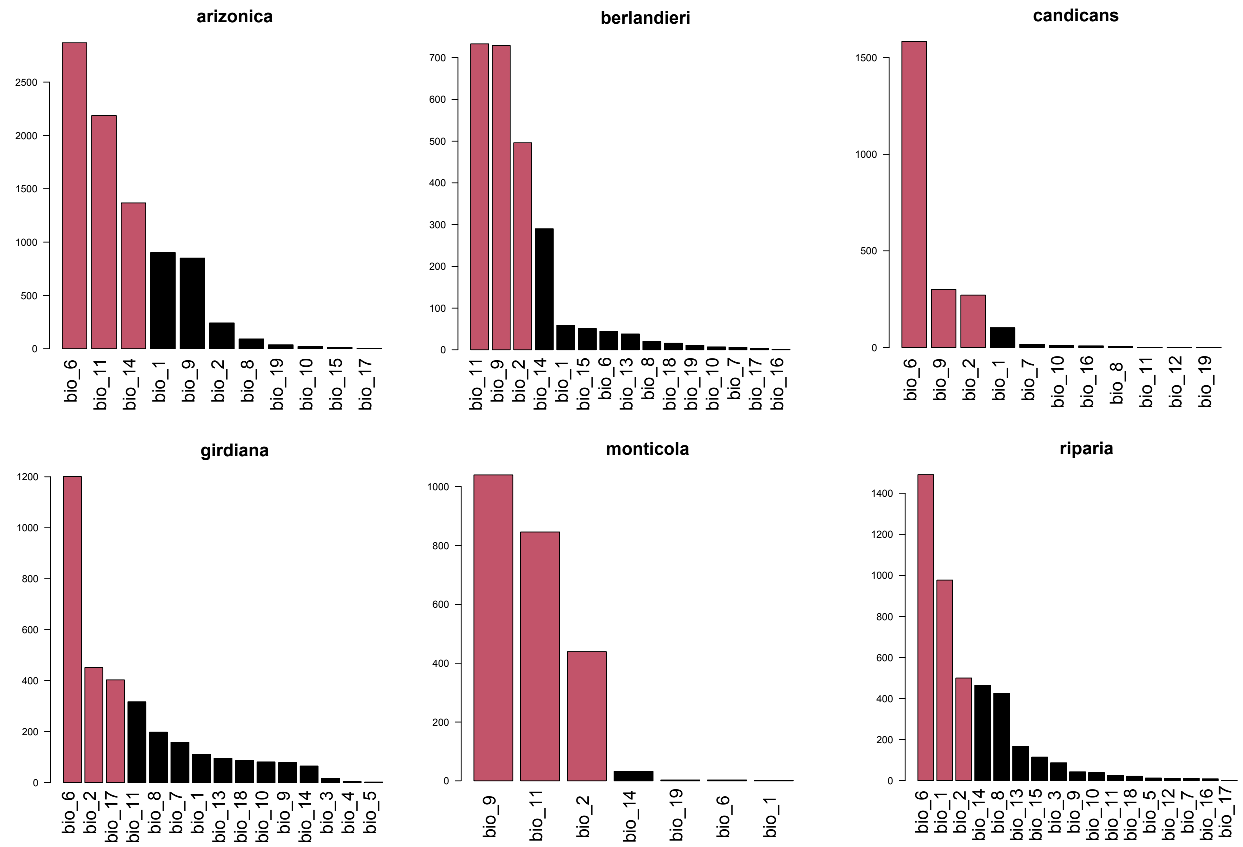


**Figure S12**. Number of associated candidate SNPs per bioclimatic variables per species. Red bars indicate the three Bioclimatic variables with the higher number of associated SNPs.

### SUPPLEMENTAL TABLES

**Table S1.** Accession identifiers and geographical coordinates of the 130 accessions used in this study. Coordinates correspond to the original collection site of the accession.

| Accession | Sample ID | Species after phylogeny | Latitude | Longitude |
|---|---|---|---|---|
| NM11-048 | vari12 | *V. arizonica* | 33.03475 | -108.20006 |
| NM11-031 | vari11 | *V. arizonica* | 32.91086 | -108.22894 |
| NM11-047 | vari21 | *V. arizonica* | 33.03644 | -108.20619 |
| NM11-040 | vari16 | *V. arizonica* | 33.03514 | -108.22894 |
| NM11-034 | vari14 | *V. arizonica* | 32.91928 | -108.21964 |
| NM11-033 | vari13 | *V. arizonica* | 32.91944 | -108.21933 |
| NM11-046 | vari07 | *V. arizonica* | 33.03928 | -108.22067 |
| NM11-035 | vari03 | *V. arizonica* | 32.92264 | -108.21567 |
| NM11-026 | vari09 | *V. arizonica* | 32.89156 | -108.23436 |
| TXNM088 | vber09 | *V. arizonica* | 29.28985 | -103.92507 |
| NM11-044f | vari20 | *V. arizonica* | 33.03994 | -108.22264 |
| NM11-044a | vari19 | *V. arizonica* | 33.03994 | -108.22264 |
| NM11-042 | vari17 | *V. arizonica* | 33.03503 | -108.22875 |
| NM11-038 | vari15 | *V. arizonica* | 32.94819 | -108.20100 |
| NM11-027 | vari10 | *V. arizonica* | 32.90675 | -108.23219 |
| NM11-039 | vari06 | *V. arizonica* | 32.94953 | -108.20667 |
| ANU58 | vrip15 | *V. arizonica* | 34.51501 | -111.76801 |
| NM11-043 | vari18 | *V. arizonica* | 33.03994 | -108.22264 |
| NM11-037 | vari05 | *V. arizonica* | 32.93492 | -108.20075 |
| NM11-036 | vari04 | *V. arizonica* | 32.92881 | -108.21033 |
| NM11-045 | vari02 | *V. arizonica* | 33.03961 | -108.22000 |
| NM11-049 | vari01 | *V. arizonica* | 33.03500 | -108.20050 |
| TX16-022 | vber21 | *V. berlandieri* | 30.84407 | -98.09657 |
| TX16-032 | vber14 | *V. berlandieri* | 30.43791 | -98.34980 |
| TX16-064 | vcan26 | *V. berlandieri* | 30.24638 | -98.04376 |
| TX16-018 | vber18 | *V. berlandieri* | 31.03509 | -98.14784 |
| TX16-065 | vber28 | *V. berlandieri* | 30.14990 | -98.05138 |
| TX16-026 | vber12 | *V. berlandieri* | 30.79163 | -98.16845 |
| TX15-003 | vber10 | *V. berlandieri* | 29.84893 | -100.02290 |
| TX16-034 | vber26 | *V. berlandieri* | 30.03024 | -98.83222 |
| TX16-035 | vber27 | *V. berlandieri* | 30.02967 | -98.83170 |

| | | | | |
|---|---|---|---|---|
| TX16-016 | vber17 | *V. berlandieri* | 31.33896 | -97.51378 |
| TX15-059 | vber11 | *V. berlandieri* | 30.43829 | -98.34963 |
| T21 | vber02 | *V. berlandieri* | 30.48940 | -99.77200 |
| T18 | vber29 | *V. berlandieri* | 30.48940 | -99.77200 |
| TX15-063 | vber23 | *V. berlandieri* | 30.68166 | -98.31197 |
| C 8-93 | vber20 | *V. berlandieri* | 31.67678 | -98.04191 |
| C 5-93 | vber19 | *V. berlandieri* | 31.16966 | -97.42910 |
| TX16-030 | vber13 | *V. berlandieri* | 30.68943 | -98.27036 |
| T 03-05 S03 | vber07 | *V. berlandieri* | 30.24687 | -98.87010 |
| TX9722 | vber05 | *V. berlandieri* | 28.70910 | -100.04950 |
| T38 | vber04 | *V. berlandieri* | 30.40020 | -97.67910 |
| T23 | vber03 | *V. berlandieri* | 30.04740 | -99.14030 |
| TX43-01 | vber01 | *V. berlandieri* | 30.31996 | -97.99865 |
| T45 | vcan02 | *V. candicans* | 30.75930 | -98.67500 |
| T64 | vcan17 | *V. candicans* | 31.13000 | -97.78000 |
| TX9715 | vcan15 | *V. candicans* | 29.34750 | -99.14140 |
| candicans 9003 | vcan29 | *V. candicans* | 32.49250 | -95.80917 |
| candicans 9005 | vcan30 | *V. candicans* | 32.49250 | -95.80917 |
| T62 | vcan16 | *V. candicans* | 31.27230 | -97.51570 |
| T36 | vcan13 | *V. candicans* | 30.40020 | -97.67570 |
| TX12-003 | vcan09 | *V. candicans* | 33.39661 | -97.50039 |
| TX32-01 | vcan04 | *V. candicans* | 30.04743 | -99.14032 |
| T48 | vcan03 | *V. candicans* | 30.75620 | -98.70030 |
| T46 | vcan14 | *V. candicans* | 30.75930 | -98.67500 |
| T2 | vcan10 | *V. candicans* | 33.25760 | -97.58324 |
| T56 | vcan08 | *V. candicans* | 31.09820 | -97.34280 |
| candicans 9039 | vcan31 | *V. candicans* | 29.91972 | -97.19306 |
| candicans 9001 | vcan28 | *V. candicans* | 31.76583 | -95.61667 |
| TX16-024 | vcan27 | *V. candicans* | 30.81022 | -98.10594 |
| TX16-006 | vcan25 | *V. candicans* | 31.10614 | -97.91810 |
| TX16-001 | vcan24 | *V. candicans* | 30.93038 | -97.52800 |
| TX14-081 | vcan23 | *V. candicans* | 30.00979 | -97.94384 |
| T 03-13 | vcan21 | *V. candicans* | 30.83118 | -99.39540 |
| T 03-08 | vcan20 | *V. candicans* | 30.16750 | -98.77183 |
| T 03-05 | vcan19 | *V. candicans* | 30.24687 | -98.87010 |
| TX9703 | vcan12 | *V. candicans* | 29.96270 | -98.78630 |
| T73 | vcan11 | *V. candicans* | 31.38660 | -97.15040 |
| SC37 | vgir19 | *V. girdiana* | 33.33007 | -117.23638 |
| NV12-040 | vgir01 | *V. girdiana* | 36.16128 | -115.49850 |
| SC11 | vgir25 | *V. girdiana* | 36.41840 | -116.31895 |

| | | | | |
|---|---|---|---|---|
| SC9 | vgir06 | *V. girdiana* | 36.41840 | -116.31895 |
| SC26 | vgir23 | *V. girdiana* | 34.93163 | -118.92795 |
| SC39 | vgir26 | *V. girdiana* | 33.31813 | -117.19465 |
| NV12-041 | vgir14 | *V. girdiana* | 36.15642 | -115.49542 |
| NV11-118 | vgir12 | *V. girdiana* | 36.37539 | -114.44308 |
| NV11-115 | vgir10 | *V. girdiana* | 36.38999 | -114.42996 |
| ANU78 | vgir02 | *V. girdiana* | 36.38925 | -114.42955 |
| NV12-050 | vgir15 | *V. girdiana* | 36.41986 | -116.32050 |
| NV11-116 | vgir11 | *V. girdiana* | 36.37678 | -114.44361 |
| SC53 | vgir17 | *V. girdiana* | 33.10527 | -117.04198 |
| SC36 | vgir27 | *V. girdiana* | 33.33007 | -117.23638 |
| SC30 | vgir22 | *V. girdiana* | 33.46235 | -117.13700 |
| SC33 | vgir21 | *V. girdiana* | 33.39145 | -117.21640 |
| SC40 | vgir20 | *V. girdiana* | 33.36293 | -117.10158 |
| SC51 | vgir16 | *V. girdiana* | 33.06660 | -116.87915 |
| C20-93-A | vmon05 | *V. monticola* | 31.06623 | -97.57890 |
| T 03-02 S01 | vmon04 | *V. monticola* | 29.98305 | -98.90348 |
| monticola 9040 | vmon07 | *V. monticola* | 32.49250 | -95.80917 |
| C20-93 | vmon06 | *V. monticola* | 31.06623 | -97.57890 |
| T40 | vmon02 | *V. monticola* | 30.30610 | -97.95240 |
| TX67-03 | vmon01 | *V. monticola* | 31.43516 | -97.74391 |
| NM12-119 | vrip21 | *V. riparia* | 36.21283 | -105.92722 |
| NM12-114 | vrip07 | *V. riparia* | 35.98822 | -105.93197 |
| NM12-108 | vrip19 | *V. riparia* | 35.32289 | -105.41953 |
| NM12-117 | vrip09 | *V. riparia* | 36.15992 | -105.97272 |
| NM12-116 | vrip08 | *V. riparia* | 35.98911 | -105.93036 |
| KS14-036 | vrip12 | *V. riparia* | 37.24091 | -99.98147 |
| CO12-103 | vrip02 | *V. riparia* | 37.11083 | -104.28486 |
| KS14-043 | vrip23 | *V. riparia* | 37.29012 | -98.61109 |
| NM12-104 | vrip18 | *V. riparia* | 35.66897 | -105.34619 |
| TXNM0824 | vrip17 | *V. riparia* | 35.66893 | -105.33610 |
| TXNM0823 | vrip16 | *V. riparia* | 35.66893 | -105.33610 |
| CO12-102 | vrip14 | *V. riparia* | 37.21572 | -104.46469 |
| OK14-064 | vrip13 | *V. riparia* | 36.53865 | -98.87695 |
| KS14-035 | vrip11 | *V. riparia* | 37.24038 | -99.98163 |
| OK14-027 | vrip10 | *V. riparia* | 36.83688 | -100.51925 |
| NM12-111 | vrip06 | *V. riparia* | 35.26753 | -105.33447 |
| V28-96 | vrip05 | *V. riparia* | 41.14871 | -73.27039 |
| V23-98 | vrip04 | *V. riparia* | 41.71946 | -72.57670 |
| OK14-026 | vrip01 | *V. riparia* | 36.83703 | -100.51897 |

| | | | | |
|---|---|---|---|---|
| T39 | vcan06 | hybrid | 30.30610 | -97.95240 |
| UT12-094 | vgir05 | hybrid | 37.29369 | -113.41414 |
| OK12-025 | vcan22 | hybrid | 34.10983 | -98.53114 |
| OK12-027 | vcan07 | hybrid | 34.11075 | -98.53250 |
| OK12-019 | vcan05 | hybrid | 34.12075 | -98.52269 |
| T42 | vcan01 | hybrid | 30.32000 | -97.99860 |
| TXNM083 | vber08 | hybrid | 30.53358 | -103.78433 |
| SC12 | vgir09 | hybrid | 37.03228 | -117.32460 |
| NV11-119 (GVC) | vgir13 | hybrid | 35.22736 | -114.68964 |
| PC96-C | vrip22 | hybrid | NA | NA |
| TX9714 | vmon03 | hybrid | 29.34750 | -99.14140 |
| SC23 | vgir24 | hybrid | 35.66705 | -118.25247 |
| SC27 | vgir08 | hybrid | 34.93163 | -118.92795 |
| SC42 | vgir07 | hybrid | 33.28781 | -116.87448 |
| UT12-084 | vgir04 | hybrid | 37.30861 | -113.42914 |
| UT12-075 | vgir03 | hybrid | 37.31033 | -113.43553 |
| candicans x aestivalis 9012 | vcan32 | hybrid | 31.76583 | -95.61667 |
| T74 | vcan18 | hybrid | 31.38660 | -97.15040 |
| T 03-01 S01 | vber06 | hybrid | 29.98305 | -98.90348 |

**Table S2.** Statistics of the *V. arizonica* b40-14 genome assembly

| Metric | Number |
|---|---|
| Total number of bases | 503291318 |
| Number of sequences | 19 |
| Average length of sequences | 2.65E+07 |
| Minimum length of sequences | 17755125 |
| Maximum length of sequences | 47005314 |
| N10 length | 35285526 |
| N20 length | 33926803 |
| N30 length | 30932466 |
| N40 length | 30043575 |
| N50 length | 25930761 |
| N60 length | 25085101 |
| N70 length | 23233325 |
| N80 length | 22502345 |
| N90 length | 19267294 |
| N10 index | 2 |
| N20 index | 3 |
| N30 index | 5 |
| N40 index | 6 |
| N50 index | 8 |
| N60 index | 10 |
| N70 index | 12 |
| N80 index | 14 |
| N90 index | 17 |
| Number of sequences >= 100bp | 19 |
| Average length of sequences >= 100bp | 2.65E+07 |
| N50 sequences >= 100bp | 25930761 |

**Table S3.** Summary table of the BUSCO genes of both haplotypes detected in the *V. arizonica* genome assembly

| Metric | Count | Percentage |
|---|---|---|
| Complete BUSCOs (C) | 1389 | 96.40% |
| Complete and single-copy BUSCOs (S) | 493 | 34.20% |
| Complete and duplicated BUSCOs (D) | 896 | 62.20% |
| Fragmented BUSCOs (F) | 14 | 1.00% |
| Missing BUSCOs (M) | 37 | 2.60% |
| Total BUSCO groups searched | 1440 | |

**Table S4.** Chromosome size, raw number of predicted SNPs and number of SNPs after filtering across all 130 samples

| Chromosome | Size (bp) | Size (Mbp) | SNPs raw bialellic | SNPs after filtering |
|---|---|---|---|---|
| Vari_b40-14_v1.hap1.chr01 | 33,926,803 | 33.9 | 2,782,419 | 1,375,689 |
| Vari_b40-14_v1.hap1.chr02 | 22,787,875 | 22.8 | 1,955,943 | 920,414 |
| Vari_b40-14_v1.hap1.chr03 | 17,755,125 | 17.8 | 1,542,440 | 767,980 |
| Vari_b40-14_v1.hap1.chr04 | 30,932,466 | 30.9 | 2,566,482 | 1,169,772 |
| Vari_b40-14_v1.hap1.chr05 | 25,930,761 | 25.9 | 2,311,363 | 1,188,958 |
| Vari_b40-14_v1.hap1.chr06 | 23,233,325 | 23.2 | 1,759,313 | 968,551 |
| Vari_b40-14_v1.hap1.chr07 | 35,285,526 | 35.3 | 2,852,943 | 1,378,311 |
| Vari_b40-14_v1.hap1.chr08 | 19,150,580 | 19.2 | 1,496,299 | 612,688 |
| Vari_b40-14_v1.hap1.chr09 | 25,085,101 | 25.1 | 2,036,641 | 931,494 |
| Vari_b40-14_v1.hap1.chr10 | 23,682,995 | 23.7 | 2,032,387 | 1,089,722 |
| Vari_b40-14_v1.hap1.chr11 | 19,267,294 | 19.3 | 1,355,229 | 665,031 |
| Vari_b40-14_v1.hap1.chr12 | 32,374,331 | 32.4 | 2,849,773 | 1,316,433 |
| Vari_b40-14_v1.hap1.chr13 | 30,043,575 | 30.0 | 2,576,413 | 1,324,160 |
| Vari_b40-14_v1.hap1.chr14 | 27,053,463 | 27.1 | 2,430,724 | 1,229,304 |
| Vari_b40-14_v1.hap1.chr15 | 22,502,345 | 22.5 | 1,999,980 | 836,821 |
| Vari_b40-14_v1.hap1.chr16 | 25,629,229 | 25.6 | 2,132,614 | 916,236 |
| Vari_b40-14_v1.hap1.chr17 | 19,376,636 | 19.4 | 1,776,932 | 918,695 |
| Vari_b40-14_v1.hap1.chr18 | 47,005,314 | 47.0 | 3,832,273 | 1,639,733 |
| Vari_b40-14_v1.hap1.chr19 | 22,268,574 | 22.3 | 1,807,138 | 741,236 |
| **Total** | 503,291,318 | 503 | 42,097,306 | 19,991,228 |

**Table S5.** Genome-wide calculation of nucleotide diversity ($\pi$) within per species

| **Species** | $\boldsymbol{\pi}$ |
|---|---|
| *V. monticola* | 0.00353 |
| *V. riparia* | 0.00312 |
| *V. candicans* | 0.00304 |
| *V. arizonica* | 0.00270 |
| *V. girdiana* | 0.00255 |
| *V. berlandieri* | 0.00211 |

**Table S6**. Results from genome-wide estimates of introgressions across all combination of trios tested.

| Trio | P1 | P2 | P3 | D | Z-score | p-value | f4-ratio | BBAA | ABBA | BABA |
|---|---|---|---|---|---|---|---|---|---|---|
| GAM | *girdiana* | *arizonica* | *monticola* | 0.086 | 15.764 | 0.00E+00 | 8.03% | 312836 | 147982 | 124492 |
| GRM | *girdiana* | *riparia* | *monticola* | 0.062 | 10.308 | 0.00E+00 | 6.08% | 294631 | 151684 | 133908 |
| ARC | *arizonica* | *riparia* | *candicans* | 0.091 | 21.944 | 0.00E+00 | 2.43% | 411536 | 126061 | 105037 |
| MBC | *monticola* | *berlandieri* | *candicans* | 0.071 | 9.224 | 0.00E+00 | 3.24% | 231272 | 189653 | 164656 |
| GRC | *girdiana* | *riparia* | *candicans* | 0.087 | 17.202 | 0.00E+00 | 2.30% | 417236 | 124661 | 104798 |
| ARB | *arizonica* | *riparia* | *berlandieri* | 0.116 | 27.752 | 0.00E+00 | 7.40% | 350794 | 140344 | 111062 |
| GRB | *girdiana* | *riparia* | *berlandieri* | 0.119 | 23.024 | 0.00E+00 | 7.47% | 357161 | 139597 | 110007 |
| AGR | *arizonica* | *girdiana* | *riparia* | 0.021 | 2.961 | 1.54E-03 | 3.32% | 204334 | 176756 | 169492 |
| RAM | *riparia* | *arizonica* | *monticola* | 0.019 | 2.737 | 3.10E-03 | 2.07% | 281783 | 151658 | 145974 |
| AGC | *arizonica* | *girdiana* | *candicans* | 0.005 | 1.256 | 1.05E-01 | 0.13% | 446358 | 106248 | 105140 |
| GAB | *girdiana* | *arizonica* | *berlandieri* | 0.001 | 0.358 | 3.60E-01 | 0.09% | 388298 | 113818 | 113480 |

**Table S7.** Gene density averages per 10 kb and relative enrichment as presented in Table 1.

| Trio | Avg. gene density | Relative. Enrichment |
|---|---|---|
| Genome-wide | 0.5762719 | - |
| ARC | 0.4417671 | 0.77 |
| GRC | 0.4383202 | 0.76 |
| GRB | 0.433835 | 0.75 |
| ARB | 0.4343598 | 0.75 |
| GAM | 0.5551546 | 0.96 |
| RAM | 0.5276498 | 0.92 |
| GRM | 0.4581006 | 0.79 |
| MBC | 0.4706927 | 0.82 |
| AGR | 0.5678322 | 0.99 |

**Table S8.** GO terms significantly enriched (p < 0.05) in pIRs

| Trio | GO ID | GO Description | p-value |
|---|---|---|---|
| AGR | GO:0005575 | C:cellular_component | 8.87E-04 |
| ARB | GO:0005575 | C:cellular_component | 5.46E-03 |
| ARB | GO:0030246 | F:carbohydrate binding | 1.81E-09 |
| GRB | GO:0030246 | F:carbohydrate binding | 9.54E-06 |
| GRC | GO:0030246 | F:carbohydrate binding | 5.34E-05 |
| GRM | GO:0030246 | F:carbohydrate binding | 5.12E-25 |
| ARB | GO:0016301 | F:kinase activity | 7.01E-07 |
| GRB | GO:0016301 | F:kinase activity | 3.82E-02 |
| GRM | GO:0016301 | F:kinase activity | 4.99E-12 |
| AGR | GO:0008289 | F:lipid binding | 9.83E-03 |
| AGR | GO:0003674 | F:molecular_function | 9.58E-03 |
| ARB | GO:0003674 | F:molecular_function | 4.59E-02 |
| ARB | GO:0000166 | F:nucleotide binding | 6.68E-07 |
| GRB | GO:0000166 | F:nucleotide binding | 2.48E-05 |
| GRC | GO:0000166 | F:nucleotide binding | 1.57E-02 |
| GRM | GO:0000166 | F:nucleotide binding | 1.07E-10 |
| ARC | GO:0004872 | F:receptor activity | 4.37E-02 |
| ARC | GO:0038023 | F:signaling receptor activity | 1.55E-02 |
| GRM | GO:0007154 | P:cell communication | 6.06E-04 |
| ARC | GO:0007267 | P:cell-cell signaling | 2.21E-07 |
| GRM | GO:0007267 | P:cell-cell signaling | 1.03E-03 |
| GRM | GO:0006464 | P:cellular protein modification process | 1.45E-05 |
| RAM | GO:0009790 | P:embryo development | 3.43E-02 |
| RAM | GO:0009908 | P:flower development | 6.48E-04 |
| GAM | GO:0006139 | P:nucleobase-containing compound metabolic process | 4.81E-02 |
| GRM | GO:0009856 | P:pollination | 5.40E-05 |

**Table S9.** Recombination rates per trio and values relative to the genome-wide estimated rate of recombination as presented in Table 1.

| Trio | Avg. cM/Kb | Relative to genome-wide |
|---|---|---|
| Genome-wide | 0.0092 | - |
| AGR | 0.0148 | 1.61 |
| ARB | 0.0166 | 1.80 |
| ARC | 0.0113 | 1.23 |
| GAM | 0.0208 | 2.27 |
| GRB | 0.0132 | 1.43 |
| GRC | 0.0134 | 1.45 |
| GRM | 0.0111 | 1.21 |
| MBC | 0.0139 | 1.52 |
| RAM | 0.0101 | 1.10 |

**Table S10.** Quantitative measurements of Pierce's Disease evaluations.

| Accession | Sample ID | Species | Category | Least Sq Mean | Std Error | Mean | Study[1] |
|---|---|---|---|---|---|---|---|
| ANU58 | vrip15 | *V. arizonica* | R | 12.485003 | 0.9391245 | 11.628 | This study |
| NM11-026 | vari09 | *V. arizonica* | R | 8.773918 | 0.9660364 | 9.424 | This study |
| NM11-027 | vari10 | *V. arizonica* | R | 10.759918 | 0.9660364 | 11.41 | This study |
| NM11-031 | vari11 | *V. arizonica* | R | 7.942103 | 0.9478225 | 7.376 | This study |
| NM11-033 | vari13 | *V. arizonica* | R | 10.741918 | 0.9660364 | 11.392 | This study |
| NM11-034 | vari14 | *V. arizonica* | R | 12.762103 | 0.9478225 | 12.196 | This study |
| NM11-035 | vari03 | *V. arizonica* | R | 10.931406 | 1.0575963 | 9.8675 | This study |
| NM11-036 | vari04 | *V. arizonica* | R | 7.707725 | 0.9528475 | 7.626 | Riaz et al 2020 |
| NM11-037 | vari05 | *V. arizonica* | R | 10.211003 | 0.6692616 | 9.354 | This study |
| NM11-038 | vari15 | *V. arizonica* | R | 8.845087 | 1.0666604 | 9.5825 | This study |
| NM11-039 | vari06 | *V. arizonica* | R | 12.227725 | 0.9528475 | 12.146 | Riaz et al 2020 |
| NM11-040 | vari16 | *V. arizonica* | R | 11.457918 | 0.9660364 | 12.108 | This study |
| NM11-042 | vari17 | *V. arizonica* | R | 11.349918 | 0.9660364 | 12 | This study |
| NM11-043 | vari18 | *V. arizonica* | R | 9.465918 | 0.9660364 | 10.116 | This study |
| NM11-044a | vari19 | *V. arizonica* | R | 9.235918 | 0.9660364 | 9.886 | This study |
| NM11-044f | vari20 | *V. arizonica* | R | 7.889725 | 0.9528475 | 7.808 | Riaz et al 2020 |
| NM11-045 | vari02 | *V. arizonica* | R | 11.181725 | 0.9528475 | 11.1 | Riaz et al 2020 |
| NM11-046 | vari07 | *V. arizonica* | R | 10.871003 | 0.9391245 | 10.014 | This study |
| NM11-047 | vari21 | *V. arizonica* | R | 11.534918 | 1.0724963 | 12.185 | This study |
| NM11-048 | vari12 | *V. arizonica* | R | 8.603603 | 1.0561199 | 8.0375 | This study |
| NM11-049 | vari01 | *V. arizonica* | R | 9.553725 | 0.9528475 | 9.472 | Riaz et al 2020 |
| TXNM088 | vber09 | *V. arizonica* | R | 10.723725 | 0.9528475 | 10.642 | Riaz et al 2020 |
| C 5-93 | vber19 | *V. berlandieri* | R | 13.102103 | 0.9478225 | 12.536 | This study |
| C 8-93 | vber20 | *V. berlandieri* | S | 14.021906 | 0.9494672 | 12.958 | This study |
| T 03-05 S03 | vber07 | *V. berlandieri* | S | 14.782248 | 0.9503933 | 14.754 | This study |
| T18 | vber29 | *V. berlandieri* | R | 11.807725 | 0.9528475 | 11.726 | This study |
| T21 | vber02 | *V. berlandieri* | R | 14.156103 | 1.2153569 | 13.59 | This study |
| T38 | vber04 | *V. berlandieri* | S | 15.851725 | 1.060632 | 15.77 | This study |
| TX15-003 | vber10 | *V. berlandieri* | S | 13.205996 | 0.9645934 | 14.906 | Riaz et al 2020 |
| TX15-059 | vber11 | *V. berlandieri* | S | 14.269906 | 0.9494672 | 13.206 | This study |
| TX15-063 | vber23 | *V. berlandieri* | S | 16.171996 | 0.9645934 | 17.872 | Riaz et al 2020 |
| TX16-016 | vber17 | *V. berlandieri* | S | 16.345996 | 0.9645934 | 18.046 | Riaz et al 2020 |
| TX16-018 | vber18 | *V. berlandieri* | S | 15.284996 | 1.0711966 | 16.985 | Riaz et al 2020 |
| TX16-022 | vber21 | *V. berlandieri* | S | 16.657496 | 1.0711966 | 18.3575 | Riaz et al 2020 |
| TX16-026 | vber12 | *V. berlandieri* | S | 17.015996 | 0.9645934 | 18.716 | Riaz et al 2020 |

| | | | | | | | |
|---|---|---|---|---|---|---|---|
| TX16-030 | vber13 | *V. berlandieri* | S | 17.053996 | 0.9645934 | 18.754 | Riaz et al 2020 |
| TX16-032 | vber14 | *V. berlandieri* | S | 15.362496 | 1.0711966 | 17.0625 | Riaz et al 2020 |
| TX16-034 | vber26 | *V. berlandieri* | S | 12.471996 | 0.9645934 | 14.172 | Riaz et al 2020 |
| TX16-035 | vber27 | *V. berlandieri* | S | 13.961996 | 0.9645934 | 15.662 | Riaz et al 2020 |
| TX16-064 | vcan26 | *V. berlandieri* | S | 14.997239 | 1.21664 | 13.9333 | This study |
| TX16-065 | vber28 | *V. berlandieri* | S | 16.885996 | 0.9645934 | 18.586 | Riaz et al 2020 |
| TX43-01 | vber01 | *V. berlandieri* | S | 16.148392 | 1.2192798 | 16.0667 | This study |
| TX9722 | vber05 | *V. berlandieri* | S | 17.903725 | 0.9528475 | 17.822 | Riaz et al 2020 |
| ANU78 | vgir02 | *V. girdiana* | R | 13.300498 | 0.8790632 | 13.6867 | Riaz et al 2020 |
| NV11-115 | vgir10 | *V. girdiana* | R | 12.263725 | 0.9528475 | 12.182 | This study |
| NV11-116 | vgir11 | *V. girdiana* | R | 10.339725 | 0.9528475 | 10.258 | This study |
| NV11-118 | vgir12 | *V. girdiana* | R | 9.753725 | 0.9528475 | 9.672 | This study |
| NV12-040 | vgir01 | *V. girdiana* | S | 16.535725 | 0.9528475 | 16.454 | This study |
| NV12-041 | vgir14 | *V. girdiana* | S | 16.669225 | 1.060632 | 16.5875 | This study |
| NV12-050 | vgir15 | *V. girdiana* | S | 16.813906 | 1.0575963 | 15.75 | This study |
| SC11 | vgir25 | *V. girdiana* | R | 14.202165 | 0.8790632 | 14.5883 | Riaz et al 2020 |
| SC26 | vgir23 | *V. girdiana* | R | 12.232165 | 0.8790632 | 12.6183 | Riaz et al 2020 |
| SC30 | vgir22 | *V. girdiana* | R | 10.857165 | 0.8790632 | 11.2433 | Riaz et al 2020 |
| SC33 | vgir21 | *V. girdiana* | S | 14.543906 | 1.21664 | 13.48 | This study |
| SC36 | vgir27 | *V. girdiana* | R | 9.368118 | 0.5385843 | 8.8462 | Riaz et al 2020 |
| SC39 | vgir26 | *V. girdiana* | R | 11.460498 | 0.8790632 | 11.8467 | Riaz et al 2020 |
| SC40 | vgir20 | *V. girdiana* | R | 14.673003 | 0.9391245 | 13.816 | This study |
| SC51 | vgir16 | *V. girdiana* | R | 11.351906 | 0.9494672 | 10.288 | This study |
| SC53 | vgir17 | *V. girdiana* | R | 10.727906 | 0.9494672 | 9.664 | This study |
| SC9 | vgir06 | *V. girdiana* | S | 16.425906 | 0.9494672 | 15.362 | This study |
| CO12-102 | vrip14 | *V. riparia* | S | 18.289725 | 0.9528475 | 18.208 | This study |
| CO12-103 | vrip02 | *V. riparia* | S | 15.681003 | 0.9391245 | 14.824 | This study |
| KS14-035 | vrip11 | *V. riparia* | S | 15.404103 | 0.9478225 | 14.838 | This study |
| KS14-036 | vrip12 | *V. riparia* | S | 15.754103 | 0.9478225 | 15.188 | This study |
| KS14-043 | vrip23 | *V. riparia* | S | 18.979003 | 0.9391245 | 18.122 | This study |
| NM12-104 | vrip18 | *V. riparia* | S | 14.587725 | 0.9528475 | 14.506 | This study |
| NM12-108 | vrip19 | *V. riparia* | S | 15.833906 | 0.9494672 | 14.77 | This study |
| NM12-111 | vrip06 | *V. riparia* | R | 14.243003 | 0.9391245 | 13.386 | This study |
| NM12-114 | vrip07 | *V. riparia* | S | 17.899003 | 0.9391245 | 17.042 | This study |
| NM12-116 | vrip08 | *V. riparia* | S | 16.71867 | 0.8586463 | 15.8617 | This study |
| NM12-117 | vrip09 | *V. riparia* | S | 17.079725 | 0.9528475 | 16.998 | This study |
| NM12-119 | vrip21 | *V. riparia* | S | 15.079725 | 0.9528475 | 14.998 | This study |
| OK14-026 | vrip01 | *V. riparia* | S | 16.903906 | 1.0575963 | 15.84 | This study |
| OK14-027 | vrip10 | *V. riparia* | S | 17.109254 | 1.2245275 | 17.8467 | This study |
| OK14-064 | vrip13 | *V. riparia* | S | 15.501906 | 0.9494672 | 14.438 | This study |

| | | | | | | | |
|---|---|---|---|---|---|---|---|
| TXNM0823 | vrip16 | *V. riparia* | R | 12.131003 | 0.9391245 | 11.274 | This study |
| TXNM0824 | vrip17 | *V. riparia* | S | 17.139003 | 0.9391245 | 16.282 | This study |
| V23-98 | vrip04 | *V. riparia* | S | 17.477104 | 1.4921791 | 16.565 | This study |
| V28-96 | vrip05 | *V. riparia* | S | 17.282104 | 1.4921791 | 16.37 | This study |

[1] The citation Riaz et al 2020 refers to Ref (Riaz et al., 2020)

**Table S11.** Number of SNPs across the genome and their association with genes for the receptor species tested in this study

| Species | PD-SNPs | N. genes with PD-SNPs | N. PD-SNPs in genes | % PD-SNPs in genes |
|---|---|---|---|---|
| *V. arizonica* | 527 | 66 | 129 | 24.5% |
| *V. berlandieri* | 5424 | 842 | 2756 | 50.8% |
| *V. girdiana* | 356 | 43 | 108 | 30.3% |
| *V. riparia* | 1690 | 208 | 625 | 37.0% |

**Table 12.** Genes with SNPs associated to PD detected in more than one species. The trio is denoted if the gene was inside pIRs.

| Chr | Gene ID | Description | PRG annotation | Assoc. Species | Trio detected |
|---|---|---|---|---|---|
| chr19 | g270290 | LRR receptor-like kinase | RLK | vber, vrip | ARB, ARC, GRB |
| chr04 | g046570 | zinc finger FYVE domain | - | vber, vrip | MBC |
| chr12 | g171820 | EI24 homolog | - | vber, vrip | ARB |
| chr12 | g167550 | Diacylglycerol kinase 5-like | CNL | vber, vgir | - |
| chr01 | g003720 | Histone chaperone ASF1B | - | vber, vrip | - |
| chr01 | g014200 | iron-sulfur cluster co-chaperone mitochondrial | - | vber, vrip | - |
| chr04 | g045340 | Ribosomal S5 domain 2-like superfamily | - | vber, vrip | - |
| chr06 | g078540 | E3 ubiquitin ligase BIG BROTHER-like | - | vber, vrip | - |
| chr08 | g115260 | eukaryotic translation initiation factor | - | vber, vgir | - |
| chr10 | g133570 | L-ascorbate oxidase homolog | - | vber, vrip | - |
| chr11 | g155590 | pentatricopeptide repeat-containing At5g52630 | - | vber, vrip | - |
| chr12 | g160220 | probable S-acyltransferase 7 | - | vari,vber | - |
| chr12 | g168300 | SNF7 family | - | vari,vber | - |
| chr14 | g196910 | nucleolar coiled-body phospho | - | vber, vrip | - |
| chr15 | g212860 | MADS-box SOC1-like | - | vber, vrip | - |
| chr15 | g213680 | probable plastid-lipid-associated chloroplastic | - | vber, vrip | - |
| chr15 | g217180 | Remorin family | - | vber, vrip | - |
| chr18 | g249790 | pesticidal crystal cry8Ba | - | vber, vrip | - |
| chr18 | g254900 | Cytochrome P450 734A1 | - | vari, vrip | - |

**Table S13.** Number of SNPs across within pIRs and their association with genes per trio

| Trio | Receptor species | PD-SNPs in pIRs | N. genes with PD-SNPs in pIRs | N. PD-SNPs in genes & pIRs | % PD-SNPs in genes & pIRs |
|---|---|---|---|---|---|
| MBC | *V. berlandieri* | 323 | 34 | 92 | 28% |
| ARB | *V. riparia* | 190 | 15 | 48 | 25% |
| GRB | *V. riparia* | 117 | 13 | 48 | 41% |
| GRM | *V. riparia* | 110 | 7 | 7 | 6% |
| GAM | *V. arizonica* | 50 | 8 | 14 | 28% |
| ARC | *V. riparia* | 18 | 3 | 6 | 33% |
| GRC | *V. riparia* | 18 | 3 | 3 | 17% |
| RAM | *V. arizonica* | 16 | 3 | 6 | 38% |
| AGR | *V. girdiana* | 0 | 0 | 0 | 0% |

**Table S14.** Comparison of number of SNPs significantly associated with PD with the expected value calculated by window permutations. P-values correspond to a t-test comparing the observed value with the whole distribution of results by random permutations.

| Trio | Observed | Expected | Relative Enrichment | p-value |
|---|---|---|---|---|
| ARB | 190 | 122.71 | 1.5484 | < 2.2e-16 |
| GAM | 50 | 39.98 | 1.2506 | < 2.2e-16 |
| MBC | 323 | 163.74 | 1.9726 | < 2.2e-16 |
| RAM | 16 | 10.16 | 1.5748 | < 2.2e-16 |
| AGR | 0 | 11.73 | 0 | 1 |
| ARC | 18 | 39.34 | 0.4575 | 1 |
| GRB | 117 | 124.81 | 0.9374 | 1 |
| GRC | 18 | 36.94 | 0.4873 | 1 |
| GRM | 100 | 100.74 | 0.9927 | 1 |

**Table S15.** Relative enrichment of the top three bioclimatic variables per trio. p-values correspond to a t-test comparing the observed value with the whole distribution of results by random permutations.

| Trio | Sp. Assoc | BIO | Relative enrich. | p-value |
|---|---|---|---|---|
| ARC | V. riparia | 1 | **1.05** | < 2.2e-16 |
| ARC | V. riparia | 2 | **1.27** | < 2.2e-16 |
| ARC | V. riparia | 6 | **1.05** | < 2.2e-16 |
| GRC | V. riparia | 1 | **1.35** | < 2.2e-16 |
| GRC | V. riparia | 2 | **1.48** | < 2.2e-16 |
| GRC | V. riparia | 6 | **1.24** | < 2.2e-16 |
| GRB | V. riparia | 1 | **1.07** | < 2.2e-16 |
| GRB | V. riparia | 2 | **1.21** | < 2.2e-16 |
| GRB | V. riparia | 6 | **1.07** | < 2.2e-16 |
| ARB | V. riparia | 1 | **1.33** | < 2.2e-16 |
| ARB | V. riparia | 2 | **1.30** | < 2.2e-16 |
| ARB | V. riparia | 6 | **1.18** | < 2.2e-16 |
| GAM | V. arizonica | 6 | 0.25 | 1 |
| GAM | V. arizonica | 11 | 0.56 | 1 |
| GAM | V. arizonica | 14 | 0.15 | 1 |
| RAM | V. arizonica | 6 | **1.34** | < 2.2e-16 |
| RAM | V. arizonica | 11 | **1.05** | < 2.2e-16 |
| RAM | V. arizonica | 14 | **1.34** | < 2.2e-16 |
| AGR | V. girdiana | 2 | **1.90** | < 2.2e-16 |
| AGR | V. girdiana | 6 | **1.07** | < 2.2e-16 |
| AGR | V. girdiana | 17 | **1.51** | < 2.2e-16 |
| GRM | V. riparia | 1 | 0.44 | 1 |
| GRM | V. riparia | 2 | 0.59 | 1 |
| GRM | V. riparia | 6 | 0.24 | 1 |
| MBC | V. berlandieri | 2 | **1.77** | < 2.2e-16 |
| MBC | V. berlandieri | 9 | **2.09** | < 2.2e-16 |
| MBC | V. berlandieri | 11 | **2.11** | < 2.2e-16 |

### DATASETS

Datasets are available is Figshare: https://doi.org/10.6084/m9.figshare.13912178

#### Dataset legends

**Dataset S1**. Genomic windows identified as putative introgressed regions (pIRs) across nine trios.

**Dataset S2**. Gene functional annotation for the genome reference of *Vitis arizonica*.

**Dataset S3**. Significant SNPs (Bonferroni adjusted $p < 0.05$) from whole genome associations with bacterial levels after infection with the causative agent of Pierce's Disease for four receptor species of introgression.

**Dataset S4**.  Significant SNPs (Bayes' Factor > 10) from whole genome associations with the top three bioclimatic variables per species.
